# AMPylation regulates PLD3 processing

**DOI:** 10.1101/2025.01.09.632087

**Authors:** Laura Hoffmann, Eva-Maria Eckl, Marleen Bérouti, Michael Pries, Aron Koller, Charlotte Guhl, Ute A. Hellmich, Veit Hornung, Wei Xiang, Lucas T. Jae, Pavel Kielkowski

## Abstract

The 5’-3’ exonuclease phospholipase D3 (PLD3) is a single-pass transmembrane protein undergoing sequential post-translational modifications (PTM) by *N*-glycosylation, AMPylation and proteolytic cleavage. The substrates of PLD3 5’-3’ exonuclease activity are single-stranded DNAs and RNAs, which act as ligands for Toll-like receptors (TLRs) and trigger a downstream pro-inflammatory response. Although PLD3 has primarily been studied in immune cells, recent findings indicate its enrichment in neurons, where it plays a role in regulating axonal fitness in Alzheimer’s disease (AD). However, the regulatory mechanisms governing the proteolytic processing of PLD3 into its catalytically active soluble form and its functional roles in both immune and neuronal cells remain unclear. Here, we describe the functional implications of PLD3 AMPylation, its direct interaction with the protein adenylyltransferase FICD, and changes in PLD3 processing in Parkinson’s disease (PD) patient-derived neurons. We identified PLD3 AMPylation sites within the proteins’ soluble region and show that mutation of these sites lead to loss of PLD3 exonuclease catalytic activity. FICD AMP-transferase accelerates PLD3 degradation and induces cellular stress response. Furthermore, depletion of the two human AMP-transferases FICD and SelO point towards a complex regulatory network governing PLD3 AMPylation. Together, our findings demonstrate a critical role of AMPylation in PLD3 processing and regulation of its catalytic activity and provide new insights into the protein’s transport and localization to lysosomes. The observation that PLD3 regulation in PD-derived neurons is altered compared to healthy neurons further highlights its role in neurodegenerative diseases.

## Introduction

In humans, two AMP-transferases, protein adenylyltransferase FICD and Selenoprotein O (SelO, gene: SELENOO) have been described to date (1–3). FICD contains a highly conserved Fic domain, which was first identified in bacteria, and is characterized by the active site sequence HPFIDGNGRTS in human cells (4, 5). The bifunctional FICD catalyzes both protein AMPylation, which involves the transfer of an adenosine monophosphate (AMP) to a target protein, and deAMPylation, the reverse process (Figure 1A). Switching between the two functions is governed by an autoinhibitory α-helix. A point mutation of the critical glutamate E234 to glycine within the autoinhibitory helix disables the enzyme’s ability to escape the highly active AMP-transferase state, which is characterized by its lack of deAMPylation activity (4, 6–11). Furthermore, the catalytic activity is regulated through FICD homodimerization and the ratio of Mg^2+^ to Ca^2+^ ions (6, 12). At the cellular level, FICD is a low abundant protein residing in the endoplasmic reticulum (ER) (13). Although FICD was initially identified as the Huntingtin-interacting protein, it has been linked to neurodegeneration in *C. elegans,* shown to accelerate neuronal differentiation in human cerebral organoids, and most recently associated with diabetes in human insulin-producing beta cells (10, 14–19).

**Figure 1.**
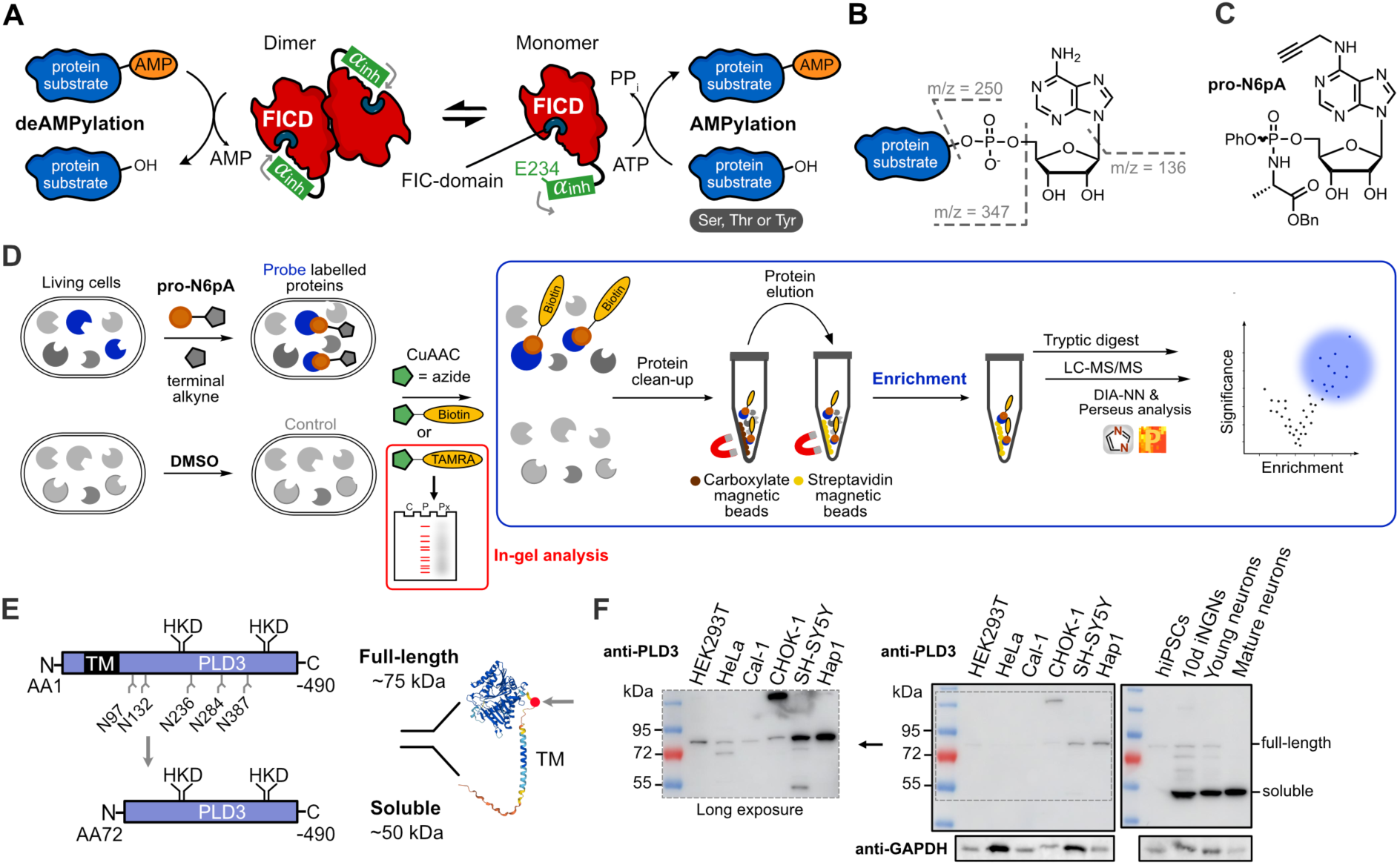
Proteomics of protein AMPylation. **A**) Schematic representation of the proposed regulation of FICD through dimerization and the autoinhibitory α-helix. **B**) Mass spectrometric properties and fragmentation of the adenosine 5’-*O*-monophosphate (AMP) group. **C**) Structure of the pro-N6pA probe. **D**) Chemical proteomics workflow used for profiling of protein AMPylation by in-gel analysis and MS-based proteomics. **E**) Schematic drawing of PLD3 domains and its proteolytic processing. AlphaFold v2.0-generated prediction of PLD3 was used for visualization (43, 44). **F**) Western blot analysis of different cell types using anti-PLD3 antibody to show differences in expression levels and proteoforms. Equal amounts (10 µg) of total protein was loaded onto SDS-PAGE. Samples were separated on two SDS-PAGEs and transferred onto one PVDF membrane for subsequent immunostaining.

The major AMPylation/deAMPylation substrate of FICD is the chaperone and unfolded protein response (UPR) regulator, heat shock protein family A member 5 (HSPA5), also known as BiP or GRP-78 (11, 13, 20). The chaperone activity of HSPA5 is inhibited by AMPylation following UPR stress release, before HSPA5 levels return to basal levels (7, 21). Another FICD AMPylation substrate has been validated *in vitro,* the peptidase Cathepsin B (CTSB), which is inhibited by AMPylation (22). However, the function and regulation of FICD-catalyzed AMPylation/deAMPylation activity and its downstream signaling role in different cell types remain poorly understood.

Direct protein AMPylation status determination by e.g. mass spectrometry-based proteomics remains challenging because of the low abundance of AMP moiety-containing peptides, poor ionization due to the presence of the negatively charged phosphate group, and the fact that AMP group fragments readily during peptide fragmentation hampering the identification in mass spectra (Figure 1B) (22–24). Several complementary strategies have been used to characterize protein AMPylation, including radioactive-labelled AMPylation substrates, anti-AMPylation antibodies and fluorescence- or affinity-tagged ATP analogues to enrich and visualize AMPylated proteins (25–29). The main drawback of ATP-based analogues is that they can be only applied in cell lysates, resulting in loss of subcellular compartmentalization. Therefore, we recently developed the *N*^6^-propagyl adenosine phosphoramidate prodrug (pro-N6pA, Figure 1C), which renders the nucleotide probe cell permeable, while avoiding the need for initial phosphorylation. This allows monitoring AMPylation in living cells (Figure 1D and Figure S1) (22, 30). Of note, the inherent competition with endogenous ATP restricts the absolute quantification of labelled proteins. The terminal alkyne on *N*^6^-propargyl is used for downstream analyses, enabling the pull-down of probe-labeled proteins by affinity purification and subsequent comparison to control or other treatment conditions.

The pro-N6pA probe has been applied to profile protein AMPylation using quantitative mass spectrometry-based chemical proteomics in human cancer cell lines and during the differentiation of human induced pluripotent stem cells (iPSCs) into neurons. This approach led to the identification of more than 100 AMPylated proteins from various subcellular localizations including the ER, cytoplasm, and lysosomes (22, 31). Significant changes in AMPylation were particularly observed on lysosomal proteins ACP2, ABHD6 and PLD3 during neuronal differentiation (31). Interestingly, neutralization of the lysosomal compartment in neuroblastoma cells by treatment with monensin or bafilomycin A1 resulted in increased AMPylation of PLD3 and the identification of an additional group of AMPylated proteins, including NOTCH2 and APP (32). Moreover, recent findings on PLD3’s genetic risk in AD, and its elevated expression levels in neurons compared to other tissues render PLD3 a possible pharmacologically target for neurodegenerative dieseases (33–37).

PLD3 is a type II transmembrane protein with a characteristic phospholipase fold, exhibiting exonuclease activity towards single-stranded DNA and RNA (Figure 1E) (38–41). Mitochondrial DNA (mtDNA) has been identified as a major physiological substrate of PLD3 in SH-SY5Y neuroblastoma cells (38). The build-up of mtDNA was caused by insufficient PINK1-dendent mitophagy in PLD3^-/-^ neuroblastoma cells (38). PLD3 is trafficked through the endolysosomal pathway to lysosomes, where it is primarily localized and undergoes proteolytical cleavage. The proteolytically cleaved soluble PLD3 has been shown to negatively regulate the immune receptor TLR9 by degrading its ligands and to positively regulate TLR7 by generating ligands for its distinct binding pockets (42). Furthermore, PLD3 is *N*-glycosylated during trafficking. The PLD3 expression levels and different proteoform ratios were shown to change dramatically during neuronal differentiation: In iPSCs, relatively low levels of full-length PLD3 form are present, while the amount of catalytically active soluble PLD3 rapidly increases in neuronal progenitors (NPCs, Figure 1F). Interestingly, during neuronal maturation, an increasing amount of soluble PLD3 becomes AMPylated, which inhibits its exonuclease catalytic activity (31).

Despite the importance of PLD3 in neurodegeneration, which may render it an interesting drug target, the mechanism and cell type-dependent differences regulating PLD3 AMPylation and its exonuclease activity in both immune cells and neurons remain unclear. Here, we established a mass spectrometry-based proteomics approach to identify protein AMPylation sites and to characterize AMPylation of PLD3 and the resulting functional consequences. We found PLD3 to directly interact with FICD and *vice versa,* suggesting that FICD is a canonical AMP-transferase modifying PLD3. Complementary proteomics analyses showed that FICD regulates PLD3 expression and processing in living cells. PLD3 processing is delayed in PD patient-derived neurons highlighting its functional role in neurodegeneration.

### Experimental Procedures

#### Culturing of HEK293T and Hap1 cells

Culturing of HEK293T and Hap1 cells HEK293T (received from Prof. Dr. Thomas Carell Lab) and Hap1 cells (received from Prof. Dr. Lucas Jae Lab) were grown in Dulbecco’s Modified Eagle’s Medium – high glucose (DMEM) or Iscove’s Modified Dulbecco’s Medium (IMDM), respectively. Both media were supplemented with 10% fetal bovine serum (FBS) and 2 mM L-alanyl-L-glutamine. Cells were maintained at 37°C in a 5% CO_2_ atmosphere. Cal-1 cells were cultured in RPMI 1640 medium supplemented with 10% heat-inactivated fetal calf serum, 100 U/ mL penicillin/streptomycin (PS), 1 mM sodium pyruvate, 2 x GlutaMAX, 10 mM HEPES and 1 x MEM NEAA. Cells were maintained in a humidified incubator at 37 °C with 5% CO_2_.

#### Cultivation of NPCs and midbrain neurons

The generation of NPCs and the differentiation to midbrain dopaminergic neurons was performed as previously described.(45) Briefly, NPCs were cultivated on 12-well plates coated with Geltrex^TM^ (Gibco) in N2B27 medium, consisting of 50% DMEM/F12 Glutamax^TM^, 50% Neurobasal Medium, 1:200 N2, 1:100 B27 and 1:100 Penicillin/Streptomycin. The N2B27 medium was further supplemented with 3µM CHIR 99021 0,5µM Purmorphamine, 10µM SB-431542 and 150µM ascorbic acid. After washing once with PBS, NPCs were split in 1:3 ratios and detached with Accutase (Sigma Aldrich). For the differentiation into midbrain dopaminergic neurons, first the medium was changed to N2B27, supplemented with 100 ng/ml FGF8, 1µM Purmorphamine and 200µM ascorbic acid for 7 days with media changes every other day. On day 8, medium was then changed to maturation medium, consisting of N2B27 medium with 10ng/mL BDNF, 10ng/mL GDNF, 1ng/mL TGF-β3, 500 μM cAMP, 0.5 μM PurMA, and 200 μM ascorbic acid. For the final replating on day 9, the maturation medium was supplemented with 10µM ROCK. On day 11, PurMA was removed from the medium, and cells were cultivated until the 16th day of maturation, with media changes every other day. iNGNs and physiological neurons were cultivated as previously described (31).

#### Probe treatment in living cells

Cells were seeded at a density of 2.5×10^6^ cells per 10 cm dish in 10 mL of culture medium. Following plating, cells were either treated with probe (100 μM pro-N6pA) or an equivalent volume of DMSO as a control. After the addition of the probe, cells were incubated for 16 hours before being harvested. Harvesting was performed by washing the cells twice with 2 mL of ice-cold DPBS, scraping them in 1 mL of DPBS, and centrifuging at 250 xg for 4 minutes at 4°C. Following centrifugation, the supernatant was removed, and the cell pellets were either frozen at −80°C or immediately subjected to cell lysis.

#### Cell lysis

Cells were lysed using 100 - 200 μL of lysis buffer (1% NP-40, 0.2% SDS, 20 mM Hepes, pH 7.5) via rod sonication (10 seconds, 20% intensity). Lysates intended for subsequent immunoprecipitation were diluted in 1 mL of lysis buffer (50 mM Tris-HCl, 150 mM NaCl, 2.5% Tween-20, pH 7.4) by shaking for 30 minutes at 4°C. The lysates were then clarified by centrifugation (13,000 xg, 4°C, 10 minutes), and the supernatant was transferred into a new 1.5 mL tube. Lysates were either stored at −80°C or processed immediately.

#### Protein concentration measurement

Protein concentration measurement was performed with a Pierce™ BCA Protein Assay Kit (Thermo Scientific).

#### Transient overexpression in human cell culture

Cells were seeded with either 2.5×10^6^ cells in 10 mL of culture medium in a 10 cm dish or 5×10^6^ cells in 15 mL for a 15 cm dish, respectively. Seeded cells were then incubated for 24 hours at 37°C with 5% CO_2_. The following day, a transfection mixture was prepared by dissolving either 10 µg or 15 µg of plasmid DNA in 1 mL or 1.5 mL of serum reduced medium (Opti-MEM^TM^). After mixing, 30 µL or 45 µL of PEI was added. The mixture was vortexed again and incubated for 20 minutes at room temperature. Meanwhile, the cell medium was replaced with fresh, pre-warmed medium before adding the transfection mixture. The cells were then incubated for another 24 hours at 37°C with 5% CO_2_. On the subsequent day, cells were either harvested as described or treated with probe/DMSO. Control samples that are not transfected are referred to as ‘untreated’ samples.

#### Depletion of FICD and SELENOO in Hap1 cells

Hap1 cell lines depleted for FICD or SELENOO were generated by utilizing the CRISPR-Cas9 system. Hap1 cells were seeded to 4×10^5^ cells per 24 well and transiently transfected 24 h later using Turbofectin (OriGene) and OptiMEM^TM^. For *FICD* the pX330 vector (Addgene 42230) containing a sgRNA (GGCGGTGACTGAACCGAAAT) targeting *FICD* exon 1 was co-transfected with a vector containing a blasticidin resistance cassette flanked by TIA sites derived from zebrafish, as described previously (46). The next day, cells were selected with 25 μg/ml blasticidin (Invivogen). Cells were selected again 4 days post transfection to ensure integration of the blasticidin resistance cassette. Clonal progeny was generated and knock-in of the blasticidin resistance cassette into the *FICD* gene was confirmed by PCR amplification of genomic DNA and parts of the resistance cassette using the following primers: 5’-GCTGCAAACAGCTAATGCACATTG-3’, 5’-GCGGGCCATTTACCGTAAGTTATG-3’, 5’-TCTTGGCTCCTTGCAGATCC-3’. Knock-in was confirmed by Sanger sequencing. SELENOO-depleted cells were constructed by co-transfection of two pX330 vectors with the sgRNAs (CCGTAGGACCTTGCGACCGT, GCCAGGCTGCCCTATACACT) targeting the second and last exon of *SELENOO*, respectively. A vector conferring transient resistance towards puromycin was co-transfected. 24 h post transfection cells were selected with 1 μg/ml puromycin. Clonal progeny was generated and genotyped by PCR using the primers 5’-GTTACACGCGTTCCCTCCTT-3’ and 5’-GGGGGAACAATGACCACAGAG-3’. genotypes were confirmed by Sanger sequencing.

#### Lentiviral transduction

Lentiviruses were generated by transfecting HEK293T cells with pFUGW_Blasticidin, containing the different transgenes, and the helper plasmids pMDLg/pRRE, pRSV-rev and pCMV-VSV-G using PEI Max. After 72 hours, the lentiviral supernatants were harvested and PLD3-deficient CAL-1 cells were transduced for 48 h and afterwards selected with blasticidin (Cat# BS.L 6115) (10 µg/mL). The polyclonal cell population was then used for further experiments.

#### Cal-1 stimulation

Cal-1 cells (100.000 cells/well in a 96-well plate) were primed with hIFN-g (10 ng/ml) for 6 hours and afterwards stimulated with 1 μg/ml R848, 5 µM CpG^S^ or 5 µM CpG^O^ DNA. Supernatants of Cal-1 cells were harvested after 16 hours of incubation at 37 °C and afterwards used to measure hIFN-b by ELISA.

#### ELISA

hIFN-b ELISA was conducted according to supplier’s protocol (Novus Biologicals).

#### Immunoblotting of PLD3 WT and PLD3 mutants expressed in Cal-1 cells

To confirm the expression of PLD3 WT and PLD3 mutants in PLD3-deficient Cal-1 cells, 1*10^6^ cells were lysed in DISC buffer (150 mM NaCl, 50 mM Tris pH 7.5, 10% glycerol, 1% Triton X-100) supplemented with cOmplete protease inhibitor cocktail for 10 minutes on ice and afterwards centrifuged for 10 minutes at 16.000 g. The supernatants were collected, 6x Laemmli buffer was added (60 mM Tris pH 6.8, 9.3% DTT (w/v), 12% SDS (w/v), 47% glycerol (v/v), 0.06% bromophenol blue (w/v)) and the samples were denatured for 5 minutes at 95 °C. Subsequently, the samples were separated by TRIS glycine SDS-PAGE and transferred onto 0.45 mm nitrocellulose membrane. Membranes were blocked in 5% milk for 1 hour at room temperature and afterwards incubated with indicated primary and corresponding secondary antibodies. Chemiluminescent signals were recorded with a CCD camera.

##### Whole proteome analysis using SP3 workflow

###### Experimental design and statistical rationale

To quantify protein levels, a full proteome analysis was conducted using DMSO-treated control lysates. Initially, 20 µg of protein from each lysate (in at least triplicates) was diluted to a total volume of 50 µL with lysis buffer. A 1:1 mixture of hydrophilic and hydrophobic carboxylate beads (Cytiva) were then equilibrated. For each sample, 20 µL of pre-mixed carboxylate beads were placed into a 96-well plate and washed three times with 100 µL of H_2_O (MS-grade). Following the final wash, the lysates were directly added to the magnetic beads. Upon addition of 60 µL of absolute ethanol, the beads were incubated for binding at room temperature for 5 minutes at 850 rpm. Subsequently, the 96-well plate was transferred to a Hamilton Microlab Prep pipetting robot to minimize pipetting errors. The plate was then placed onto a magnet to remove supernatant containing unbound components. The magnetic beads were washed thrice with 80% ethanol in H_2_O (MS-grade) and once with acetonitrile (ACN). For each wash, the beads were incubated for 1 minute at room temperature while shaking at 850 rpm in the washing solution, followed by placement of the plate on the magnet to remove the wash solution. Finally, the beads were dissolved in 60 µL of 100 mM ABC buffer, and 1 µL of sequencing-grade trypsin (0.5 mg/mL, Promega) was added for overnight digestion at 37°C while shaking at 650 rpm. The following day, the 96-well plate was returned to the magnet, and the supernatant containing the peptide mixture was transferred to a clean 1.5 mL tube. The beads were manually washed twice with 50 µL and 30 µL of 1% FA, respectively. Each time, the plate was incubated for 5 minutes at 40°C while shaking at 850 rpm. The supernatants were combined in the initial 1.5 mL tube, which was then reattached to the magnet. Finally, the supernatant was transferred to an MS-vial and subjected to LC-MS/MS analysis. Unless mentioned otherwise, whole proteome samples were prepared in triplicates derived from one cell lysate.

##### Enrichment of modified proteins using small-scale SP2E workflow

###### Experimental design and statistical rationale

The procedure was carried out using the Hamilton Microlab Prep pipetting robot in a 96-well plate format (with at least duplicates for each condition). Initially, 100 μg of in-cellulo probe-labelled (or DMSO control) lysates were diluted to 40 μL with lysis buffer. CuAAC-mediated click reaction including 0.4 μL of biotin-N_3_ (10 mM in DMSO) and 0.25 μL of TBTA (16.7 mM in DMSO) were then added. Additionally, either 0.4 μL of TCEP (Condition A; 100 mM in MS-H_2_O) or 1.2 µL TCEP (Condition B; 100 mM MS-H_2_O) were added. Condition A was applied to experiments shown in figures 2A, 2D, 2E, 6E and 6F. Condition B was applied to experiments shown in figures 2C, 5C, 5D, 6E, 6H. The reaction was initiated by adding 0.8 μL of CuSO_4_ (50 mM in MS-H_2_O) and incubating at room temperature, 600 rpm for 1.5 hours. Following incubation, 40 μL of urea (8 M in MS-H_2_O) was added to the click reaction to reach a final volume of 80 μL. A 1:1 mixture of hydrophobic and hydrophilic carboxylate-coated magnetic beads (Cytiva) was washed three times with 100 μL of MS-H_2_O. The reaction mixture was then transferred onto 20 µL of the pre-washed beads and all further steps were carried out by the pipetting robot. The workflow starts with the addition of 100 μL absolute ethanol (EtOH), mixing, and incubation at room temperature, 850 rpm for 5 minutes. Subsequently, the beads were washed three times with 150 μL of 80% EtOH in MS-H_2_O and once with 150 μL MS-grade ACN. Proteins were eluted three times with 60 μL of 0.2% SDS in PBS at 40 °C, 850 rpm for 5 minutes from the carboxylate-coated beads and transferred onto 50 μL pre-equilibrated streptavidin-coated magnetic beads (New England Biolabs) by a 3x wash with 100 μL of 0.2% SDS in PBS. For streptavidin-biotin complex formation, beads were incubated at room temperature, 800 rpm for 1 hour with eluted proteins. Subsequently, beads were washed three times with 150 μL of 0.1% NP-40 in PBS, two times with 150 μL of 6 M urea (in MS-H_2_O), and two times with 150 μL of MS-H_2_O by incubating at room temperature, 800 rpm for 1 minute between each wash step. After the last wash step, 50 μL of TEAB buffer (50 mM in MS-H_2_O) was added, and digestion was carried out overnight with 1.5 μL sequencing-grade trypsin (0.5 mg/mL, Promega) at 37 °C, 600 rpm. The next day, supernatants were transferred into new 1.5 mL tubes, and beads were washed manually once with 20 μL of TEAB buffer (50 mM in MS-H_2_O) and once with 20 μL of 0.5% MS-grade formic acid (in MS-H_2_O) by incubating each time at 40 °C, 600 rpm for 5 minutes. To the combined fractions, 0.9 μL MS-grade formic acid was added, placed onto the magnet for 15 minutes, and transferred into new 1.5 mL tubes. These samples were placed onto the magnet for 2 minutes, the supernatant was transferred into MS vials to ensure that no beads were in the samples and subjected to LC-MS/MS measurement. If not mentioned otherwise, enrichment samples were prepared in triplicates derived from one cell lysate.

**Figure 2.**
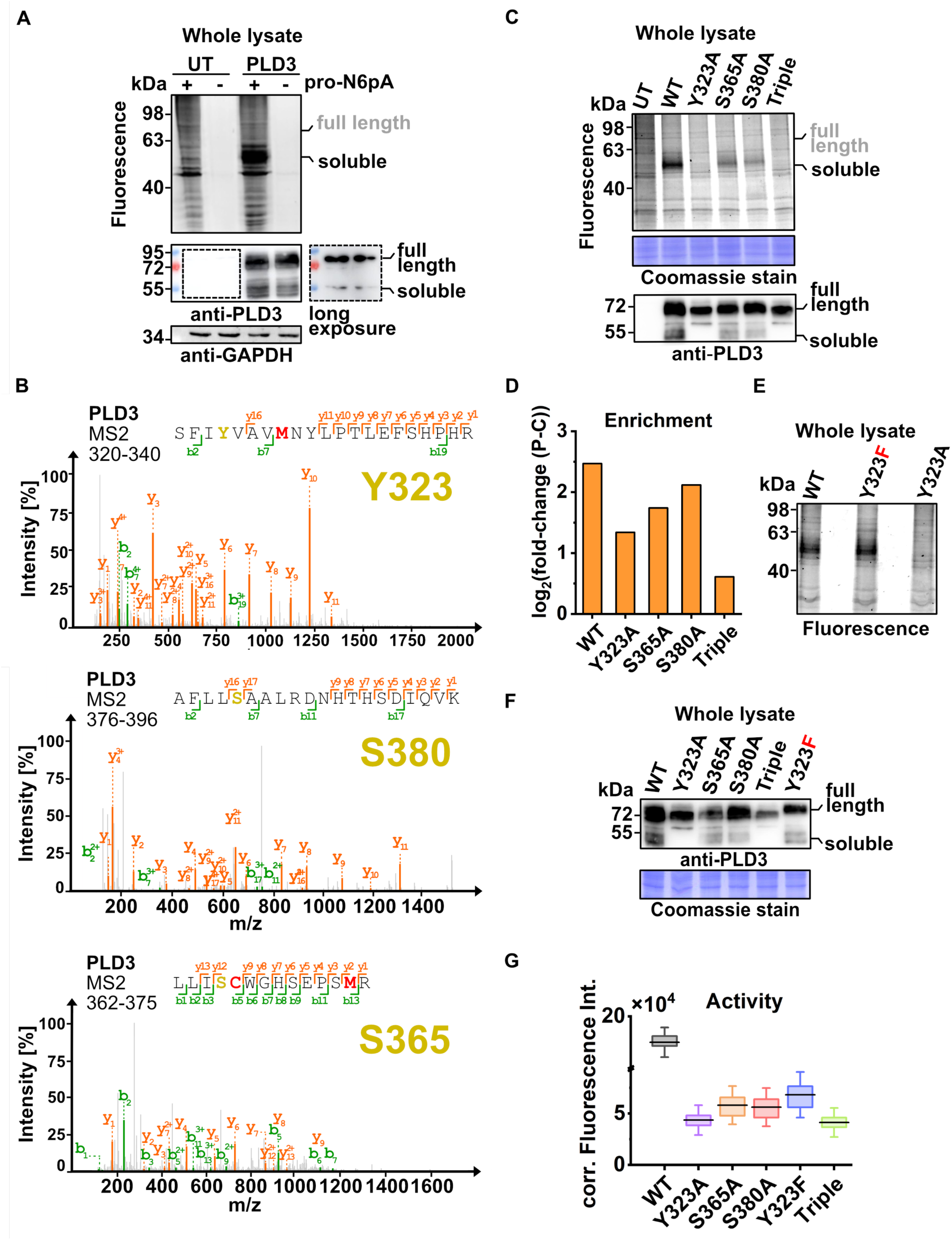
Identification and validation of PLD3 AMPylation sites. **A**) HEK293T cells transiently transfected with a vector encoding full-length PLD3 subsequently treated with pro-N6pA and visualized with TAMRA-azide via CuAAC. The figure shows a fluorescence scan and Western blot of untreated (UT) control and overexpressed PLD3 in HEK293T cells. The strong increase of fluorescence signal at around 55 kDa indicates that mostly the soluble form of PLD3 is AMPylated. One representative blot of three independent experiments is shown. **B**) Representative MS^2^ spectra of newly identified AMPylation sites Y323 (HR, DIA, 10 ppm), S380 (standard, DIA, 10 ppm) and S365 (HR, DIA, 10 ppm). In total, sites were identified in 8 spectra for S365 (four containing natural AMP and four with N^6^-propargyl AMP) and two independent replicates; eight spectra for Y323 (eight containing natural AMP) in three independent replicates; two spectra for S380 (two natural AMP) and three independent replicates (Figures S3). **C**) Fluorescence scan of PLD3 single point mutants Y323A, S365A and S380A as well as the triple mutant. Strong or complete depletion of pro-N6pA probe labelling for all three single and and the triple mutant corroborated validates these three as the main AMPylation. The Western blot of PLD3 shows that Y323A and triple mutant fails to produce the soluble PLD3 form. One representative blot of two independent experiments is shown. **D**) AMPylation profiling indicates the decrease in AMPylation of PLD3 variants (Figure S6). **E**) Fluorescence-based analysis of the PLD3 Y323F point mutant shows that in this mutant, AMPylation on the other sites is retained. One representative gel of two independent experiments is shown. **F**) PLD3 Y323F retains formation of the soluble form but without catalytic activity (see Figure 2E). One representative blot of two independent experiments is shown. **G**) Box plot visualizing the decrease in PLD3 exonuclease catalytic activity, *n* = 3.

##### Enrichment of modified proteins using large-scale SP2E workflow

###### Experimental design and statistical rationale

400 μg of proteins from in-cellulo control or probe-treated lysates were diluted to 200 μL with lysis buffer (each condition prepared in at least duplicates). CuAAC-mediated click reagents including 2 μL of biotin-PEG-N_3_ (10 mM in DMSO), 6 μL of TCEP (100 mM in MS-H_2_O), and 0.25 μL of TBTA (83.5 mM in DMSO) were then added, vortexed, and briefly spun. The reaction was initiated by adding 4 μL of CuSO_4_ (50 mM in MS-H_2_O) and incubating at room temperature, 600 rpm for 1.5 hours. After completion of the incubation, 200 μL of urea (8 M in MS-H_2_O) was added to the click reaction to reach a final volume of 400 μL. A 1:1-mixture of hydrophobic and hydrophilic carboxylate-coated magnetic beads (Cytiva) was washed three times with 500 μL of MS-water, and the reaction mixture was then transferred onto the 50 µL per sample of pre-washed beads, followed by the addition of 600 μL absolute ethanol (EtOH), mixing, and incubation at room temperature, 950 rpm for 5 minutes. The beads were washed three times with 500 μL of 80% EtOH in MS-water. Proteins were eluted twice with 500 μL of 0.2% SDS in PBS at room temperature, 950 rpm for 5 minutes from the carboxylate-coated beads and transferred onto 50 μL pre-equilibrated streptavidin-coated magnetic beads (New England Biolabs) by a 3x wash with 1 mL of 0.2% SDS in PBS. For streptavidin-biotin complex formation, beads were incubated at room temperature, 950 rpm for 1 hour with eluted proteins. Subsequently, beads were washed three times with 500 μL of 0.1% NP-40 in PBS, two times with 500 μL of 6 M urea (in MS-H_2_O), and two times with 500 μL of MS-H_2_O by mixing and briefly spinning between each wash step. After the last wash step, 25 µL of 5x Laemmli buffer (10% (w/v) SDS, 50% (v/v) glycerol, 25% (v/v) β-mercaptoethanol, 0.5% (w/v) bromphenol blue, 315 mM Tris/HCl, 100 mM DTT, pH 6.8) was added and incubated at 95°C for 5 minutes. Supernatants were transferred into new Eppendorf tubes and either stored at −80°C or directly loaded onto an SDS-Gel for subsequent western Blot analysis. If not mentioned otherwise, enrichment samples were prepared in one replicate derived from one cell lysate.

##### PLD3 exonuclease activity assay

###### Experimental design and statistical rationale

For quantitative measurement of PLD3 5’ exonuclease activity, lysates were prepared in TRIS-lysis buffer (TBS with 1% (v/v) Triton X-100 and 1 tablet of protease inhibitor). After collecting the whole cell lysates as described above, lysates were diluted in triplicates to a final volume of 100 µL in MES reaction buffer (50 mM MES, 200 mM NaCl, pH 5.5). The final concentration was adjusted to 50 ng/µL for untreated cells, 1 ng/µL for wt PLD3 and 5 ng/mL for PLD3 point mutants in a lumox multiwell 96 plate (Sarstedt). Each sample was measured in triplicates, *n=3*. The reaction was initiated by adding 100 pmol of quenched FAM-ssDNA substrate (6-FAM-ACCATGACGTTC*C*T*G*-BMN-Q535 from Biomers.net, with * indicating a phosphothioate bond). Fluorescence emission at 528 nm (excitation at 485 nm) was measured using a microwell plate reader (Tecan) over a 4-hour period, with measurements taken every 5 minutes at 37°C. For evaluation, a substrate control without lysate and a lysate control without substrate were measured alongside the samples. The corrected fluorescence intensity, Fc(t), was calculated as the mean fluorescence intensity for each time point minus the values from the two controls.

###### Immunoprecipitation using anti-FLAG affinity gel

For the immunoprecipitation of overexpressed FLAG-tagged protein, anti-FLAG® M2 Affinity Gel (Thermo Fisher) was used. The gel was equilibrated according to the manufacturer’s protocol (Part II: Resin Preparation). The immunoprecipitation procedure was carried out as described in the section FLAG® Fusion Protein Immunoprecipitation of the manual. All centrifugation steps were carried out at 4°C for 1 min and 6000 xg. Briefly, 40 μL of gel suspension per reaction (equivalent to 20 μL of packed gel volume) were used. As controls, either a reagent blank with no protein or protein lacking the FLAG-tag (negative control) was employed. Triplicates were prepared for each condition: duplicates for LC-MS/MS analysis and one replicate for Western blot analysis. For each reaction, 1 mg of in-cellulo probe-treated (or DMSO control) lysates were added to the equilibrated gel matrix and the volume was adjusted to 1 mL with lysis buffer (50 mM Tris-HCl, 150 mM NaCl, Tween-20, pH 7.4). The gel was incubated for 2 hours at 4°C with shaking to facilitate protein binding to the matrix. After incubation, the supernatant was removed, and the gel was washed twice with 0.5 mL of wash buffer (50 mM Tris-HCl, 150 mM NaCl) and twice with MS-grade water (MS-H_2_O). Next, an in-gel CuAAC click reaction was performed. To the gel, 1 µL of TAMRA-N_3_ (10 mM in DMSO), 1 µL TCEP (100 mM in H_2_O), and 0.125 µL TBTA (83.5 mM in DMSO) were added, and the mixture was adjusted to a final volume of 100 µL with 1x PBS. The reaction was initiated by adding 2 µL CuSO_4_ (50 mM in H_2_O) and incubated for 1.5 hours at room temperature with mixing at 650 rpm. Samples intended for Western blot analysis were centrifuged at room temperature for 1 minute at 6000 xg, the supernatant was discarded, and the proteins were eluted from the gel by adding 25 µL of 1x Laemmli buffer and heating for 5 minutes at 95°C and 850 rpm. The supernatants were transferred to fresh tubes and loaded onto the SDS-Gel. Samples destined for LC-MS/MS analysis were centrifuged, the supernatant was discarded, and 80 µL ABC buffer (125 mM in MS-H_2_O) and 10 µL TCEP (100 mM in MS-H_2_O) were added. The sample was heated to 56°C for 30 minutes for reduction. Afterwards, 10 µL CAA (400 mM in MS-H_2_O) was added, and the mixture was incubated for 30 minutes at room temperature for alkylation. For digestion, 1.5 µL of sequencing-grade Trypsin (Promega) was added, and the samples were incubated overnight at 37°C. The next day, samples were vortexed and spun down for 30 seconds at room temperature at 6000 xg. Trypsin activity was quenched by acidification using 2 µL of 100% MS-grade formic acid. Samples were desalted using 50 mg SepPak C18 columns. The columns were equilibrated by adding 1 mL of 100% ACN, followed by 1 mL 80% ACN/0.5% formic acid in MS-H_2_O, and finally three times with 1 mL 0.5% fromic acid in MS-H_2_O. The sample was then loaded and washed three times with 1 mL 0.5% formic acid in MS-H_2_O (gravity flow). The column was removed, the aperture was dried by applying a vacuum, and finally, the proteins were eluted into fresh 1.5 mL tubes by adding 250 µL 80% ACN/0.5% formic acid in MS-H_2_O (gravity flow) followed by 250 µL 80% ACN/0.5% formic acid in MS-H_2_O (vacuum). The peptide mixture was dried in the SpeedVac for 2.5 hours at 35°C, and the dried peptide mixture was dissolved in 30 µL 1% formic acid before transfer into MS vials. Samples were analyzed using short and long HPLC gradients combined with high-resolution MS and MS/MS measurement and the DIA method. This experiment was reproducibly conducted at two different time points, preparing biological triplicates (three individual cell dishes) for each condition (pro-N^6^pA probe or DMSO control) on each occasion. Duplicates were utilized for LC-MS/MS measurements, while one replicate was allocated for SDS-Gel analysis. Additionally, the duplicates intended for LC-MS/MS analysis were measured three times.

###### Endoglycosidase H-digest

For cleaving N-linked glycosylation of glycoproteins, recombinant Endoglycosidase H (NEB) was employed. 20 μg of glycoprotein, 1 μl of 10X Glycoprotein Denaturing Buffer, and H_2_O were combined to create a total reaction volume of 10 μl. Glycoproteins were denatured by heating the reaction at 100°C for 10 minutes. Subsequently, 2 µL of 10X GlycoBuffer 3 and 1 µL of EndoH were added, and the mixture was filled up to 20 µL with H_2_O. The reaction mixture was then incubated for 1 hour at 37°C.

###### Chemical Transformation of Bacterial cells

For chemical transformation, chemically competent E. coli DH5-alpha cells stored at −80°C were thawed for 10 minutes on ice. Then, 50 ng of plasmid DNA or 5 µL of the KLD reaction mixture (see site-directed mutagenesis section) was added directly to 50 µL of cells and the tube was gently flicked three times. The cells were incubated for 30 minutes on ice and then exposed to 42°C for 30 seconds, a process known as heat shock. The cells were immediately placed on ice for 5 minutes for regeneration before the addition of 950 µL of SOC medium (NEB) and incubation for 1 hour at 37°C at 180 rpm. In the meantime, LB-Agar (1.3% Agar) plates supplemented with 100 µg/mL ampicillin were pre-warmed at 37°C. After incubation, 100 µL of the cell suspension was spread onto pre-warmed LB-Agar plates and evenly distributed using glass beads by shaking the plate under sterile conditions. The LB-Agar plates were then incubated overnight at 37°C and subsequently stored at 4°C or directly used for plasmid amplification.

###### Plasmid DNA isolation using Mini/Midi Preparation Kit

For plasmid DNA isolation, either the Mini (small scale) or Midi (large scale) preparation kit from NEB was utilized. For Mini or Midi preparation, 3 mL or 100 mL, respectively, of sterile LB medium supplemented with 100 µg/mL ampicillin was prepared. A single clone from the overnight LB agar plate was selected using a pipette tip and subsequently added to the medium. The medium was then incubated for 12-16 hours at 37°C at 180 rpm. Plasmid DNA was isolated following the manufacturer’s protocol for the Plasmid Miniprep/Midiprep Kit (NEB). The DNA was eluted in deionized H_2_O. DNA concentration was measured using a Nanodrop, and the DNA sequence was confirmed by Sanger sequencing conducted by Genewiz.

###### Q5-site-directed mutagenesis

For introduction of point mutations to plasmids, the Q5-site directed mutagenesis kit was used. To validate the AMPylation sites in PLD3, point mutants of the appropriate sites were prepared. For this purpose, a primer pair containing the mutated amino acid for each of the three potential AMPylation sites (Y323, S365, S380) was designed. The triple mutant containing all three mutations was prepared in an iterative manner. Here, purified DNA template of the first point mutant previously confirmed by sequencing was used for the introduction of the second- and third-point mutation by repeating the described steps with different primer pairs. The forward and reverse oligonucleotides as well as the applied annealing temperature (T_a_) for cloning the different PLD3 construct are listed below in **Error! Reference source not found.**S1. The soluble form of PLD3 was cloned starting from amino acid position 61 and adding an N-terminal IgK-signal peptide using the full-length wildtype PLD3 with a C-terminal FLAG(3x)-tag as a template. For cloning the catalytically inactive point mutant FICD H363A, the following primer pair was used together with the template containing wildtype FICD. For introduction of C-terminal FLAG(3x)-tag to FICD wildtype or mutants (H363A and E234G) a primer pair containing the FLAG(3x)-tag sequence (5’-DYKDHDGDYKDHDIDYKDDDDK-3’) was prepared. For all constructs, the sequence identity was validated by Sanger sequencing (Genewiz) using the appropriate sequencing oligonucleotides listed in **Error! Reference source not found.**S2.

###### Polymerase chain reaction (PCR)

PCR was performed for cloning purposes using plasmid DNA as a template. Here, procedure was performed according to manufacturers’ protocol (Q5 site-directed mutagenesis; NEB). The thermocycling conditions applied are listed below in Table S3. PCR product formation was validated by 1% Agarose gel electrophoresis.

###### Kinase, Ligase and DpnI (KLD) reaction

After PCR, KLD reaction was performed for circularization of the linear product based on manufacturers’ protocol (Q5 site-directed mutagenesis; NEB). Here, the PCR product is first phosphorylated (Kinase), then ligated (Ligase) and residual template DNA is decomposed by DpnI enzyme (only methylated DNA is a substrate). The reaction mixture was prepared, mixed by pipetting up and down 5 to 10 times and incubated for 5 min at room temperature. The KLD mixture was used for subsequent bacterial transformation.

###### 1%-Agarose Gel

For validation of PCR product formation, 1% agarose gel electrophoresis was conducted. The agarose was dissolved in 40 mL 1x TAE buffer (40 mM Tris, 20 mM acetic acid, 1 mM EDTA) and heated up in the microwave. After cooling down, 2 µL Gelstain was added and the gel was polymerized. As a standard, 1kB DNA ladder (Carl Roth) was used. To the PCR product 6x stain (NEB) was added (c_f_ = 1x). The electrophoresis was conducted for 1h at 100 V in 0.5x TAE buffer. PCR products were detected under UV light.

###### Western Blot analysis

For Western blot analysis, 20 µg of cell lysate was used. Proteins were denatured using 5x Laemmli buffer (10% (w/v) SDS, 50% (v/v) glycerol, 25% (v/v) β-mercaptoethanol, 0.5% (w/v) bromphenol blue, 315 mM Tris/HCl, 100 mM DTT, pH 6.8) and diluted to a final concentration of 1x. Then, samples were boiled for 5 min at 95°C and subsequently loaded onto a 10% SDS-Gel thereby allowing protein separation by size. The separated proteins were transferred onto a polyvinylidene fluoride (PVDF) membrane in a semi-dry manner. For this purpose, the sodium dodecyl sulfate (SDS)-gel and filter paper was equilibrated in transfer buffer (48 mM Tris, 39 mM glycine, 0.0375% (m/v) SDS, 20% (v/v) methanol) for 5 min at room temperature. The PVDF membrane was first activated for 1 min in MeOH and then equilibrated in transfer buffer for 5 min at room temperature. To allow transfer, the filter paper was placed on the bottom with the membrane on top followed by the SDS-gel and another filter paper on top. Then, the protein transfer was carried out for 30 min at 25 V using a semi dry blotter (Bio-Rad). Afterwards, non-specific binding sites were blocked by incubating the membrane for 1h at room temperature in blocking solution (0.5 g milk powder in 10 mL PBST (PBS/0.5% (v/v) Tween)). Subsequently, 10 mL of primary antibody diluted in blocking solution with specificity for the protein of interest was added and the mixture was incubated for 1 hour up to overnight at 4°C. The membrane was washed 3 times for 10 min with PBST before 1 mL of the secondary antibody diluted in 10 mL of blocking solution was added. As loading control fluorophore-coupled anti-GAPDH antibody was added during the secondary antibody incubation. After 1 hour of incubation at room temperature in the dark, the membrane was washed three times for 10 min with PBST while shaking at room temperature. Then, a 1:1 mixture composed of 400 µL of ECL substrate and 400 µL peroxide solution was prepared and added to the membrane to for chemiluminescence signal detection. Finally, images of the western blot were taken by developing using the Amersham Imager 680 (GE Healthcare) machine.

###### Lysosomal immunoprecipitation (LysoIP)

For isolation of the lysosomal fraction using anti-HA beads, lysosomal membrane protein TRPML1 with HA-tag was overexpressed in HEK293T cells. For each condition, two 150 cm dishes were seeded, each containing HEK293T cells. Cells were incubated for 24 hours at 37°C and 5% CO_2._ The next day, cells were transfected using 15 µg plasmid DNA containing TRPML1 gene with 3xHA tag and again incubated for 24 hours at 37°C and 5% CO_2_. As a control, untreated (not transfected) HEK293T cells were used. The next day, cells were washed twice with ice-cold PBS and collected by scraping in 7 mL KPBS I buffer (10 mM KH_2_PO_4_, 136 mM KCl, pH 7.25, freshly added protease inhibitor tablet) per plate into a fresh 15 mL falcon. The falcon was spun down for 5 min at 200 xg and 4°C. The supernatant was discarded, and the pellet was resuspended in 1 mL KPBS I buffer. Then, 80 µL of magnetic anti-HA beads per condition were equilibrated thrice with KPBS I buffer by pipetting up and down. Beads were resuspended in 80 µL KPBS I buffer. Cell suspension was transferred to a tissue grinder (VWR) to ensure cell lysis by preserving organellar membrane. For this, cells were homogenized by 30x strokes thereby avoiding air bubble formation. Homogenate was transferred to fresh 1.5 mL tubes. An aliquot of 20 µL was used as ‘whole cell lysate (L)’ fraction and diluted with 20 µL lysis buffer and 5 µL 5x Laemmli for subsequent SDS-PAGE analysis. The homogenate was centrifuged for 2 min at 4°C and 2000 xg and the supernatant containing lysosomal fraction was transferred directly to equilibrated beads and resuspended by pipetting twice up and down. To allow binding, beads were incubated for 10 min at 4°C under continuous motion. The tubes were placed on a magnet and the supernatant containing the ‘flow through (FT)’ fraction was removed after taking an aliquot of 20 µL and diluting with 20 µL lysis buffer and 5 µL 5x Laemmli. Next, beads were washed twice with 500 µL KPBS III buffer (100 mM KH_2_PO_4_, 25 mM KCl, 150 mM NaCl pH 7.2, freshly added protease inhibitor tablet) and once with KPBS II buffer (100 mM KH_2_PO_4_, 25 mM KCl pH 7.2, freshly added protease inhibitor tablet). After each washing step, the mixture was transferred to a fresh 1.5 mL tube to avoid contamination of unbound fraction. Finally, the last washing solution was aspirated and the proteins were eluted from the beads with 120 µL elution buffer (KPBS I Buffer, 0.5% NP-40, freshly added protease inhibitor tablet). Elution occurred while incubating beads at 4°C for 30 min while continuous motion. Tubes were placed on the magnet and supernatant was transferred to fresh 1.5 mL tube and spun down for 10 min at 4°C and 13.000 xg. The Supernatant was transferred to fresh 1.5 mL tubes and an 40 µL aliquot of ‘immunoprecipitation (IP)’ fraction was diluted with 10 µL 5x Laemmli for subsequent SDS-PAGE analysis. Eluates of LysoIP were stored after snap-freezing in liquid N_2_ at −80°C. Samples for SDS-PAGE analysis were heated up to 95°C for 5 min and 15 µL of each fraction was loaded onto a 10% SDS-Gel followed by western blot analysis. For whole proteome analysis via LC-MS/MS, protein concentration of eluates was measured via BCA assay and continued with SP3 workflow (as described). The experiment was conducted at two different time points, preparing biological duplicates (two individual cell dishes) for TRPML1-HA overexpressing cells and one replicate for untreated control cells on each occasion. LysoIP samples were prepared in duplicates from each lysate for LC-MS/MS measurements. One replicate was allocated for SDS-Gel analysis.

###### In-gel fluorescence analysis

For in-gel fluorescence analysis, 100 μg of in cellulo probe-labelled or DMSO-control lysates were diluted to 100 μL with lysis buffer (1% NP-40, 0.2% SDS, 20 mM Hepes, pH 7.5). Click reagents such as 1 μL of TAMRA-N_3_ (10 mM in DMSO), 3 μL of TCEP (100 mM in MS-water) and 0.125 μL of TBTA (83.5 mM in DMSO) were added, vortexed and spun down. The reaction was initiated by the addition of 2 μL of CuSO_4_ (50 mM in MS-water) and the mix was incubated at room temperature while shaking at 600 rpm for 1.5 hours. Proteins were precipitated with 400 μL pre-cooled acetone (4:1 ratio) at −20 °C for at least 1 hour and were pelletized for 10 min at 4 °C and 13,000 xg. Protein pellet was reconstituted in 80 μL 0.2% SDS in PBS. 20 μg total protein lysate per sample was boiled for 5 min at 95 °C with 5x Laemmli buffer, loaded and run with 150 V for 1 hour in 1x running buffer (25 mM Tris, 0.192 M glycine, 0.1% (m/v) SDS) on a 10% SDS-PAGE gel to electrophoretically separate the proteins. In-gel fluorescence was scanned with Amersham Imager 680 (GE Healthcare). For loading control, proteins were stained with a Coomassie staining solution (0.25% Coomassie Blue R-250, 10% acetic acid, 50% MeOH) and properly destained with destaining solution (20% MeOH, 10% acetic acid).

###### LC-MS/MS measurement

MS measurements were conducted using an Orbitrap Eclipse Tribrid Mass Spectrometer (Thermo Fisher Scientific) connected to an UltiMate 3000 Nano-HPLC (Thermo Fisher Scientific) via a Nanospray Flex (Thermo Fisher Scientific) and FAIMS interface (Thermo Fisher Scientific). Peptides were initially loaded onto an Acclaim PepMap 100 μ-precolumn cartridge (5 μm, 100 Å; 300 μM ID x 5 mm, Thermo Fisher Scientific), followed by separation at 40 °C on a PicoTip emitter (noncoated, 15 cm, 75 μm ID, 8 μm tip, New Objective) packed in-house with Reprosil-Pur 120 C18-AQ material (1.9 μm, 150 Å, Dr. A. Maisch GmbH). LC buffers comprised MS-grade water (A) and acetonitrile (B), both supplemented with 0.1% formic acid. The short gradient ran from 4-35.2% B over 60 minutes (0-5 min: 4%, 5-6 min: 7%, 7-36 min: 24.8%, 37-41 min: 35.2%, 42-46 min: 80%, 47-60 min: 4%) at a flow rate of 300 nL/min. The long gradient spanned 150 minutes, with a B gradient from 4-35.2% (0-5 min: 4%, 5-6 min: 7%, 7-105 min: 24.8%, 105-126 min: 35.2%, 126-140 min: 80%, 140-150 min: 4%) at a flow rate of 300 nL/min.

###### Data-independent acquisition

Field asymmetric ion mobility spectrometry (FAIMS) was conducted with a single CV set at −45 V. Each DIA cycle consisted of one MS^1^ scan followed by 30 MS^2^ scans. The mass spectrometer operated in DIA mode with the following parameters: Polarity: positive; MS^1^ Orbitrap resolution: 60k for standard or 120k for HR; MS^1^ AGC target: standard; MS^1^ maximum injection time: 50 ms; MS^1^ scan range: m/z 200-1800 or if mentioned m/z 120-1700; RF Lens: 30%; Precursor Mass Range: m/z 500-740; isolation window: m/z 4; window overlap: m/z 2; MS^2^ Orbitrap resolution: 30k or 60k for HR; MS^2^ AGC target: 200%; MS^2^ maximum injection time: auto; HCD collision energy: 35%; RF Lens: 30%; MS^2^ scan range: auto.

###### Data-dependent acquisition

FAIMS was performed with two alternating CV settings, −50 V and −70 V. The mass spectrometer operated in dd-MS^2^ mode with the following parameters: Polarity: positive; MS^1^ Orbitrap resolution: 240k; MS^1^ AGC target: standard; MS^1^ maximum injection time: 50 ms; MS^1^ scan range: m/z 375-1500; RF Lens: 30%; MS^2^ Orbitrap resolution: 15k; MS^2^ AGC target: standard; MS^2^ maximum injection time: 35 ms; HCD collision energy: 30%; RF Lens: 30%; MS^2^ cycle time: 1.7 s; intensity threshold: 1.0e4 counts; included charge states: 2-6; dynamic exclusion: 60 s.

###### DIA-NN

First, *.raw files were converted to *.mzML format using “MSConvert” from the “ProteoWizard” software package (http://www.proteowizard.org/download.html) with the following settings: “peakPicking” filter with “vendor msLevel = 1”; “Demultiplex” filter with parameters “Overlap Only” and “mass error” set to 10 ppm. Subsequently, they were analysed with DIA-NN 1.8.1, and peptides were searched against the UniProt database for *Homo sapiens* (taxon identifier: 9606) with included contaminants and decoys. The DIA-NN settings were as follows: FASTA digest for library-free search/library generation: enabled; Deep learning-based spectra, RTs, and IMs prediction: enabled; missed cleavages: 2; max number of variable modifications: 3; modifications: N-term M excision, carbamidomethylation, oxidation (M), and N-term acetylation; Precursor charge range: 2-6; precursor range: m/z 500-740; fragment ion range: m/z 200-1800; precursor FDR level: 1%; match between runs (MBR): enabled; library generation: smart profiling; quantification strategy: Robust LC; mass accuracy: 0; scan windows: 0.

###### FragPipe

MS *.raw files were converted to *.mzML format with “MSConvert” with following settings: “peakPicking” filter with “vendor msLevel = 1”, “titleMaker” filter was enabled. Peptides were searched in Fragpipe 1.8 against Uniprot database for Homo sapiens (taxon identifier: 9606) with included contaminants and decoys. For protein digestion, trypsin (cuts at KR) was chosen with 2 missed cleavages. Due to the huge variety of settings, general workflows provided by FragPipe were used and subsequently modified accordingly. All results were viewed using the integrated FP-PDV viewer and directly from the psm.tsv results table. Results in the psm.tsv file were filtered for being ‘unique’ and for q-value ‘<0.1%’. In the following most important features were listed for both, AMPylation and *N*-Glycosylation site-identification. only hits fulfilling these parameters were taken into consideration.

###### AMPylation site identification using MSFragger-DIA search engine

For AMPylation site-identification, general workflows for DIA data, DIA_SpecLib_Quant and DIA_SpecLib-Umpire, offered by FagPipe were used and adjusted according to **Error! Reference source not found.**S4.

###### N-Glycosylation site identification using MSFragger-Glyco search engine

Glycosylation site identification was performed using template glycol workflow provided by MSFragger-Glyco. The following two basic workflows were applied and successfully results in detection of N-Glycosylation sites: *glyco-N-HCD* and *glyco-N-open-HCD* (open search), both for CID/HCD fragmentation of N-glycopeptides. Glycoproteomics results were viewed using the integrated FP-PDV viewer and directly from the psm.tsv results table.

###### Volcano plot preparation from quantified data (DIA-NN)

For statistical analysis the “report.pg_matrix.tsv” table was used in Perseus 1.6.14.0.21 Then, quantified values were log_2_-transformed and columns were assigned with either control- or probe-treated. Subsequently, the groups were filtered based on the experimental background (at least two valid values out of three columns in at least one group or at least two valid values out of three columns in each group). Further, missing values were replaced from a normal distribution. - log_10_(*p*-values) were obtained by a two-sided one sample Student’s t-test over replicates with the initial significance level α = 0.05 adjustment by the multiple testing correction method of Benjamini and Hochberg (FDR = 0.05) using the volcano plot function. Finally, the volcano plot values from Perseus software (1.6.10.43) were transferred to OriginPro 2022 (9.9.5.171) and visualized properly.

###### Principal component analysis (PCA)

For Principal component Analysis, quantified values were log_2_-transformed, and columns were assigned to the cell lines tested (Hap1 WT, Hap1 FICD KO, or Hap1 SELENOO KO) using Perseus software (1.6.10.43). Subsequently, the groups were filtered, and missing values were replaced based on normal distribution. PCA was conducted using the integrated tool in Origin 2022b (9.9.5.171).

###### Statistical analysis

All statistical analyses were conducted using Origin 2022b (9.9.5.171). For multiple group comparisons against each other, a one-way ANOVA with Tukey’s range test was performed, also correcting for multiple comparisons to a significance level of α ≤ 0.05. For comparisons between only two groups, a two-sided t-test was employed with a significance level of α ≤ 0.05.

## Results

### PLD3 processing is cell type dependent

We previously showed that in neurons, the soluble form is the major PLD3 proteoform (31). Here, we compared the expression levels and proteoforms of PLD3 in several cell types (Figure 1F). We observed that in cancer cell lines and plasmacytoid dendritic cells (Cal-1), the full-length, membrane-bound PLD3 is indeed the main form, while in neuronal progenitors (NPCs) and neurons the pool of PLD3 is mainly composed of proteolytically cleaved soluble PLD3 (Figure 1F). Quantification of the total PLD3 levels revealed that on average 17-fold more PLD3 is present in neurons in comparison to other tested cell types (Figure S2). Interestingly, the highest level of full-length PLD3 in non-neuronal cells was determined in Hap1 cells, a near haploid cell line derived from chronic myeloid leukemia (47).

### Multiple sites within the soluble region of PLD3 are AMPylated

Identifying AMPylation sites on endogenous proteins in living cells remains a significant challenge. Thus, it remains unclear whether the lack of reported AMPylation sites on PLD3 is due to their absence, which seemed unlikely due to the importance of this PTM, or the technical challenges associated with detecting such sites, such as low abundancy of soluble AMPylated PLD3 in the cell lines used, leading to a small pool of modified peptides to be detected. Additionally, the poor ionization efficiency of phosphate group-containing modified peptides and the lability of *N*-glycosidic bond during fragmentation contribute to the challenge (23, 24, 26). To increase the amount of putatively AMPylated PLD3 peptides for MS analysis, we linked full-length PLD3 to a C-terminal 3xFLAG-tag (PLD3-FLAG) enabling efficient immunoprecipitation (IP) of the protein from whole cell lysates. PLD3-FLAG was transiently overexpressed in HEK293T cells which were subsequently treated with either pro-N6pA or DMSO as a control. Subsequent Western blot analysis revealed high expression levels of PLD3-FLAG protein. In-gel fluorescence analysis, utilizing the CuAAC reaction with TAMRA-azide to specifically tagged AMPylated proteins in whole cell lysates, revealed a significant increase in the fluorescent band around 55 kDa, corresponding to luminal soluble PLD3. Lysates from non-transfected cells, referred to as untreated (UT), were compared to PLD3 overexpressing cells. A strong fluorescence signal was observed, particularly in the lysates from probe-treated cells, in contrast to the DMSO-treated control (Figure 2A). This points towards correct processing of PLD3 and successful incorporation of pro-N6pA into PLD3. Together, the Western blot and in-gel fluorescence analysis showed a notable increase in PLD3 AMPylation confirming the feasibility of our approach.

Next, overexpressed PLD3 was immunoprecipitated using anti-FLAG agarose beads followed by on beads-digest with trypsin for MS analysis. Digested samples were analysed by different LC-MS/MS methods on Orbitrap Tribrid Eclipse using short (35 min) and long (90 min) LC gradients coupled to MS analysis running in data independent acquisition (DIA) mode or data dependent acquisition (DDA) mode combining standard high resolution orbitrap for MS^1^ and faster low-resolution linear ion trap for MS^2^. While the DDA approach did not revealed desired AMPylation sites, analysis of DIA data with MSFragger-DIA(48, 49) spectral search with natural AMPylation (+329.0525) or *N*^6^-propargyl AMP (+367.0682) on Ser, Thr and Tyr as variable modification revealed multiple AMPylation sites on Y323, S365 and S380 of PLD3 with high confidence (Figure 2B and Figure S3). Natural AMP and the *N*^6^-propargyl AMP group could be detected at all sites showing even distribution of our AMP homolog in the target protein. While the *N*^6^-propargyl AMP was not found in DMSO treated controls, with the exception of a single poor-quality spectrum for S365, which can be assigned as false positive due to known challenging false discovery rate (FDR) estimation for modified peptides and complexity of MS^2^ spectra in DIA mode (50). To further confirm the identified AMPylation sites, newly prepared PLD3 IP samples were remeasured using DIA method with higher-resolution of MS^2^. With this approach we confirmed that all sites contain AMP. Interestingly, residue S380 was recently shown to be involved in homodimeric PLD3 interactions and to be located within the dimer interface suggesting that the AMPylation status may modulate PLD3 catalytic activity (40). Furthermore, the Y323 site is in spatial proximity to a transmembrane domain suggesting a potential role in regulation of PLD3 proteolytic cleavage (40).

To validate the identified AMPylation sites, PLD3 point mutants were generated in which the respective Ser or Tyr were replaced by alanine residue. The PLD3 point mutants were subsequently overexpressed in HEK293T cells and incubated with pro-N6pA to determine the AMPylation status by in-gel fluorescence. For PLD3 Y323A, the fluorescent signal based on pro-N6pA was completely depleted (Figure 2C). For both, the S365A and S380A mutant, a strong decrease in fluorescent signal was observed pointing towards lack of AMPylation on these sites (Figure 2C). The complementary Western blot analysis of the PLD3 forms revealed absence of the luminal PLD3 for Y323A and the triple mutant containing all three point mutations, and somewhat lower levels for mutants S365A and S380A. The total PLD3 levels of all point mutants were comparable with overexpressed wildtype (wt) PLD3 as shown by Western blot and confirmed by proteomics (Figure 2C and Figure S4). Interestingly, while there was no major change in the whole proteome of cells expressing the single PLD3 point mutants, the expression of the triple PLD3 mutant exhibited strong dysregulation of whole proteome, with over 500 proteins being significantly upregulated and 160 downregulated (Figure S4). The GO term search points towards changes in RNA metabolic processes, RNA processing and ribonucleoprotein complex biogenesis (Figure S5). The overall decrease of AMPylation on PLD3 was confirmed by pull-down of AMPylated proteins using the pro-N6pA probe, coupled via CuAAC with biotin, followed by affinity enrichment and LC-MS/MS analysis (SP2E, Figure 2D, Figure S6 and Table S5). Of note, HEK293T cells also contain endogenous PLD3.

The unexpected complete depletion of PLD3 AMPylation on all identified sites in the single point-mutant Y323A led us to construct an additional point mutant exchanging Y323 to structurally more similar phenylalanine. In this mutant, AMPylation at residues S365 and S380 could be partially restored (Figure 2E). The proteolytic processing of the PLD3 Y323F mutant was restored as well, as observed by Western blot analysis (Figure 2F). Taken together, the described MS-based experiments provide direct confirmation of PLD3 AMPylation and identify three modified amino acid side chains Y323, S365, and S380 within the soluble domain of PLD3.

### AMPylation is necessary for PLD3 catalytic activity in HEK293T cells

Next, we asked how the point mutations of the AMPylation sites influence the PLD3 5’-3’ exonuclease activity. A previously described PLD3 activity assay was utilized, which relies on the cleavage of a short ssDNA substrate releasing fluorescent signal upon scission (41). The activity assay was conducted in cell lysates overexpressing either wt PLD3 or its point mutants. The catalytic activity of PLD3 Y323A and triple mutant were completely inhibited, while the other PLD3 variants including S365A, S380A and Y323F displayed significantly decreased activities compared to the WT enzyme (Figure 2G and Table S6). We speculate that the PLD3 Y323A mutation may cause improper folding and proteolytic processing of the full-length protein, resulting in a lack of the active soluble form.

In human cancer cell lines, immune Cal-1 cells, and iPSCs, the major PLD3 form is the full-length protein, whereas in neuronal progenitor cells, the soluble form is more abundant. Finally, in mature neurons, the main form is AMPylated PLD3 leading to inhibition of its catalytic activity (Figure 1F, Figure S7B and S7D) (31). This suggests a crucial role for PLD3 AMPylation and processing for neuronal development. Comparing PLD3 exonuclease activity across different cell lines indicate that the amount of soluble non-AMPylated PLD3 is crucial for its catalytic activity (Figure S7A). Furthermore, based on the activity data of PLD3 Y323A, Y323F, S365A and S380A mutants, we propose that PLD3 AMPylation is a necessary transient modification leading to release of the soluble active PLD3, which in neurons might be halted at the stage of AMPylated soluble form to block its exonuclease activity (Figure S7B, S7C, and S7D). The soluble active PLD3 might be then swiftly released by deAMPylation from relatively large pool of inactive enzyme. Together these data show that AMPylation is necessary to establish the catalytic activity of PLD3.

### Depletion of PLD3 AMPylation leads to TLR9 activation in Cal-1 cells

To assess whether PLD3 AMPylation mutants lack the ability to process single-stranded DNA in immune cells we made use of the pDC-like immortalized cell line Cal-1. It was previously described that PLD3 degrades DNase 2-generated ligands for the Toll-like receptor TLR9 in mice (34). Consequently, it was shown that CpG^O^-DNA did not trigger a TLR9 response in wildtype Cal-1 cells but in the absence of PLD3, indicating that PLD3 also negatively regulates the TLR9 response in Cal-1 cells (42). We therefore stably expressed wt PLD3 and PLD3 AMPylation mutants in PLD3-deficient Cal-1 cells (Figure 3A) and stimulated those cells with CpG^O^-DNA as well as TLR agonist R848 as a control (Figure 3B and 3C, Figure S7E). As expected, wildtype Cal-1 cells did not respond to CpG^O^-DNA stimulation, while CpG^O^ stimulation in PLD3-deficient cells induced robust IFN-β release (Figure 3B). Overexpression of the wt PLD3 protein in PLD3-deficient cells reversed this effect. PLD3 S365A and PLD3 S380A overexpression also suppressed the CpG^O^ response, similar to the wt PLD3 protein, indicating that PLD3 S365A and PLD3 S380A are still functional in degrading DNA in Cal-1 cells. Interestingly, overexpression of PLD3 Y232A and the triple mutant PLD3 (S365A, S380A, Y323A) and subsequent stimulation with CpG^O^ resulted in partial IFN-β release for PLD3 Y232A and robust IFN release for PLD3 (S365A, S380A, Y323A), mimicking the PLD3-deficient cell line. TLR agonist R848 control stimulation resulted in similar IFN-β release in all conditions tested (Figure 3C). Stable expression of mCherry in PLD3-deficient Cal-1 cells was further used as a control and showed similar behavior to the PLD3-deficient cell line. Furthermore, PLD3 S365A, S380A, Y323A mutant resulted in somewhat smaller changes on whole proteome compared to overexpression in HEK293T cells (Figure 3D, Figure S4 and Table S7). Together, in Cal-1 pDC-like immune cells, dysregulation of PLD3 AMPylation leads to aberrant TLR9 responses in PLD3 Y323A and triple mutant containing cells.

**Figure 3.**
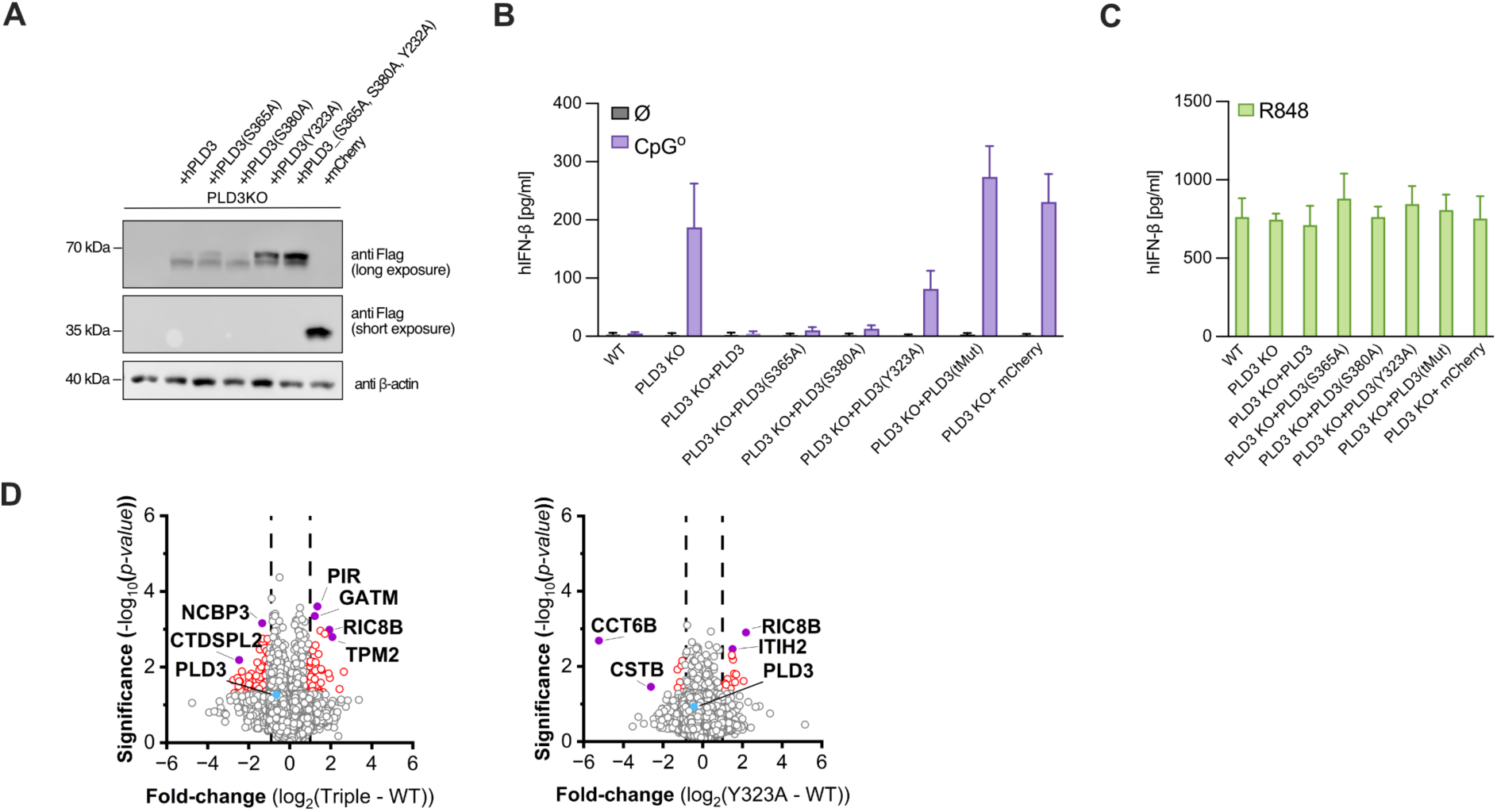
PLD3 dependent TLR9 activity in CAL-1 cells. **A**) Western blot of wt and *PLD3^-/-^* cells reconstituted with PLD3-FLAG, indicated PLD3-FLAG mutants or mCherry (FLAG). One representative blot of three independent experiments is shown. **B**) wt Cal-1, *PLD3^-/-^* or *PLD3^-/-^* cells reconstituted with wt PLD3, indicated PLD3 mutants or mCherry were unstimulated or stimulated with CpG^O^ and IFN-β release was measured by ELISA. Data are depicted as mean ± SEM of n = 3 independent experiments. **C**) wt Cal-1, *PLD3^-/-^* or *PLD3^-/-^* cells reconstituted with wt PLD3, indicated PLD3 mutants or mCherry were unstimulated or stimulated with R848 and IFN-β release was measured by ELISA. Data are depicted as mean ± SEM of n = 3 independent experiments. **D**) Whole proteome analysis of Cal-1 cells expressing PLD3 point mutants (see also Figure S8).

### The AMP-transferase FICD directly interacts with PLD3

To elucidate whether the endoplasmic reticulum resident AMP-transferase FICD interacts with PLD3, we utilized the PLD3-FLAG construct to co-immunoprecipitate (co-IP) interacting proteins. To account for unspecific background binding proteins, we compared the PLD3 co-IP from HEK293T cells transfected without the affinity tag to those transfected with the PLD3-FLAG construct. Since the levels of endogenous FICD are low and difficult to detect by both, Western blot and MS-based proteomics, in parallel to PLD3 transient overexpression, the wt FICD containing vector was used to increase the FICD levels. First, to assess the efficiency of the PLD3-FLAG IP, Western blotting using anti-PLD3 antibody was carried out and showed an efficient pull-down of PLD3-FLAG from the lysate (Figure 4A), while in the control condition without the FLAG-tag, PLD3 was not enriched (Figure 4A and Figure S11). The following Western blot with the anti-FICD antibody revealed the corresponding FICD band, which was missing in the co-IP containing PLD3 construct without the FLAG tag (Figure 4B). To corroborate the FICD-PLD3 interaction, the wt FICD-FLAG construct was used to transiently overexpress FICD and wt PLD3 (without an affinity tag). As expected, the co-IP of FICD-FLAG showed a clear enrichment of PLD3 (Figure 4B). Interestingly, only one intermediate form of PLD3 was observed, which was neither corresponding to the ∼72 kDa full-length form nor to the soluble PLD3 form around 55 kDa. The soluble PLD3 is known to be stabilized by *N*-glycosylation; in the mouse protein on residues N97, N132, N234, N282 and N385 (39, 40).

**Figure 4.**
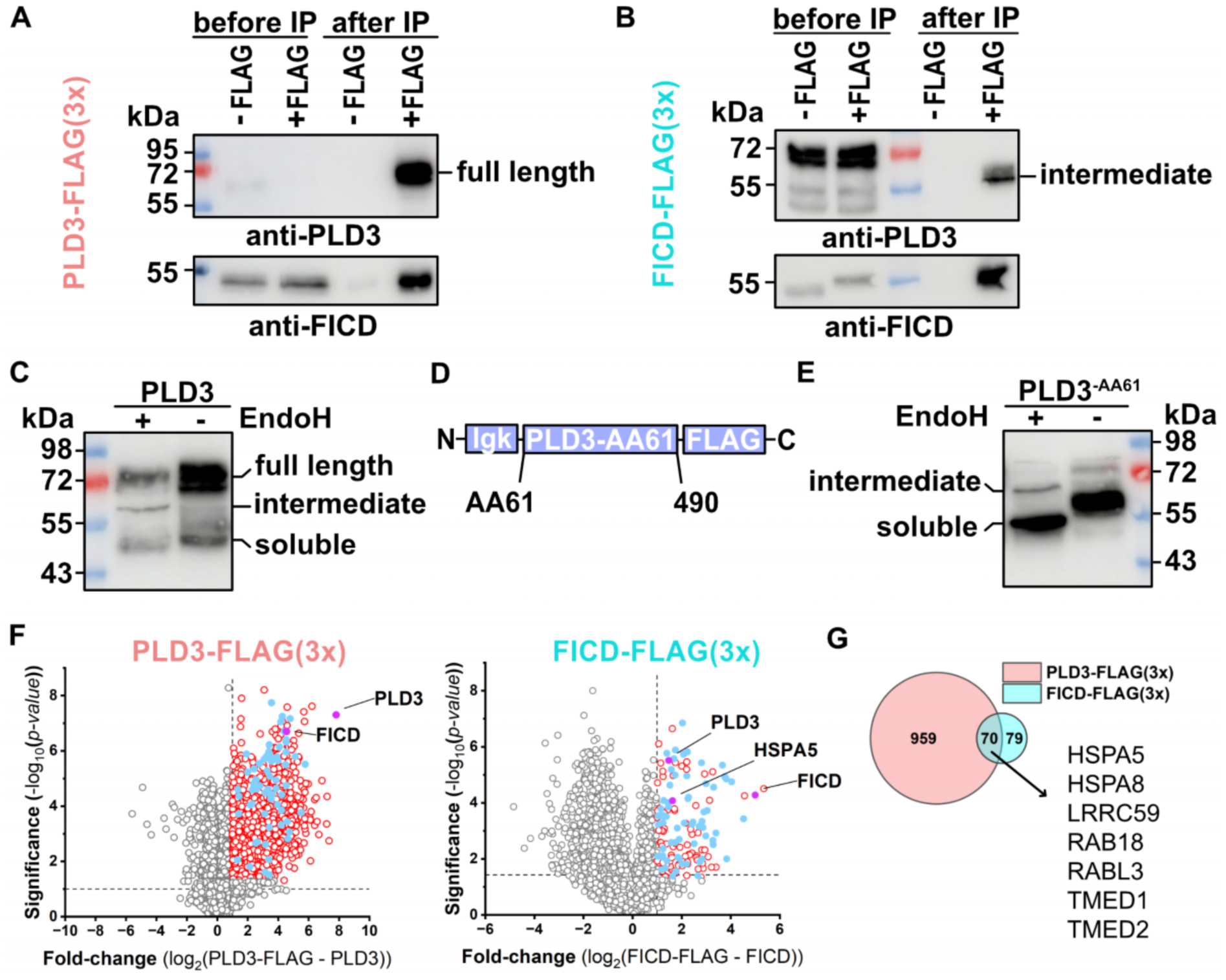
FICD interacts with the intermediate form of PLD3 in HEK293T cells. **A**) Western blot analysis before and after immunoprecipitation of PLD3-FLAG using anti-FLAG beads demonstrates co-immunoprecipitation of FICD. One representative blot of two independent experiments is shown. **B**) Western blot analysis before and after immunoprecipitation of FICD-FLAG using anti-FLAG beads for cross-validation confirms PLD3 and FICD interaction and further elucidates soluble, glycosylated PLD3 interacting with FICD. **C**) Western Blot analysis of PLD3-FLAG before and after incubation with Endo H glycosidase specifically cleaving *N*-linked glycoproteins showing almost quantitative *N*-glycosylation of full-length PLD3. **D**) Truncated luminal PLD3^-AA61^ equipped with *N*-terminal ER targeting sequence. **E**) Western Blot analysis of PLD3^-AA61^ before and after incubation with Endo H glycosidase showing almost quantitative *N*-glycosylation of full-length PLD3. **F**) Volcano plots visualizing the results of co-IP MS-based proteomics of PLD3-FLAG and FICD-FLAG constructs. Red circles – significantly enriched proteins, blue dots – proteins found significantly enriched in both PLD3 and FICD co-IPs, gray circles – not significantly enriched proteins. Co-IPs were prepared in triplicates (*n = 3)* from one protein lysate. **G**) Venn diagram showing the overlap between significantly enriched proteins in PLD3 and FICD co-IPs.

To confirm that the *N*-glycosylation takes place in our expression system, the MS data from PLD3 immunoprecipitation were searched by MSFragger for *N*-glycosylation sites to corroborate previously identified N236 and N387 glycosylation sites on human PLD3 (51, 52). Furthermore, in order to distinguish the individual PLD3 forms separated by SDS-PAGE followed by in-gel analysis or Western blotting, the lysates were incubated with Endo H glycosidase specifically cleaving *N*-linked glycoproteins. The comparison of the bands observed before and after the Endo H incubation showed almost quantitative *N*-glycosylation of full-length PLD3 (Figure 4C). Next, the interaction of FICD and PLD3 was corroborated by MS-based proteomics (Figure 4F and Table S8) to show an overlap of 70 proteins between the two IPs (Figure 4G). Together, these data point towards direct interaction between FICD, PLD3 and HSPA5 suggesting a crosstalk between UPR and TLR9 signaling.

### Hap1 FICD-depleted cells corroborate cell type specificity of PLD3 AMPylation

To further investigate the interaction between FICD and PLD3, we targeted FICD with single-guide RNAs (sgRNAs) in near haploid Hap1 cells. The sgFICD line was complemented by an sgSELENOO line to analyze both thus far two known AMP-transferases. First, we determined the changes on whole proteome level, which were induced by depletion of FICD or SELENOO (Figure 5A and Table S9). Principal component analysis (PCA) revealed distinct and significant changes in protein expression (Figure 5B). More than 300 proteins were downregulated in both cell lines. The most downregulated proteins, including SKP2 and FBXL6, are involved in G2/M transition of mitotic cells cycle and regulating protein localization to the membrane such as TCAF1, GPC3 and GPC4. Interestingly, heat-shock protein HSPB1 was one of most downregulated proteins in the sgFICD cells and also significantly downregulated in sgSELENOO cells (Figure 5A).

**Figure 5.**
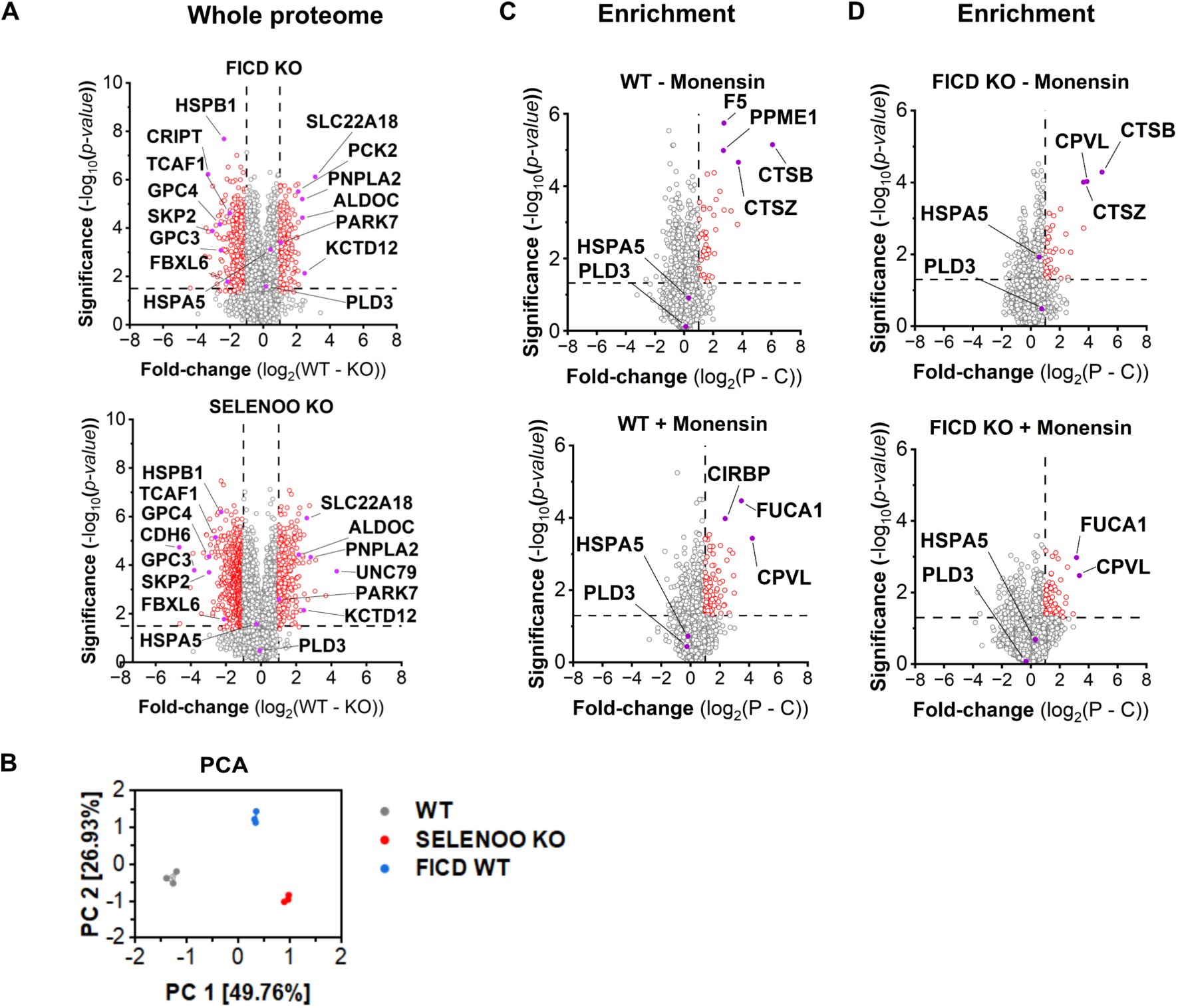
Absence of PLD3 AMPylation in HAP1 cells underscores cell-type specificity of this PTM?. **A**) Whole proteome changes in FICD wt / KO and SELENOO wt/KO cells. Representative result from two independent measurements. Samples were prepared in triplicates (*n = 3)* from one protein lysate. **B**) PCA analysis for wt, SELENOO KO, and FICD KO cells. **C**) Enrichment analysis of wt cells with (+) or without (-) Monensin treatment. Samples were prepared in triplicates (*n = 3)* from one cell lysate. **D**) Enrichment analysis of FICD KO cells (+) or without (-) Monensin treatment. Samples were prepared in triplicates (*n = 3)* from one protein lysate.

In contrast, HSPB1 was found upregulated in cells overexpressing FICD E234G, see below. Furthermore, the 85 proteins upregulated in both sgFICD and sgSELENOO cells include ALDOC, PCK2, PNPLA2, SLC22A18, KCTD12 and PARK7 pointing towards the role in carbohydrate metabolism and oxidative stress regulation (Figure 5A).

Next, we analyzed the overall AMPylation profile in Hap1 utilizing the pro-N6pA probe for enrichment followed by LC-MS/MS analysis (Figure 5C and Table S10). However, there was no enrichment observed neither for HSPA5 nor PLD3, while other previously identified AMPylated proteins were found such as PPME1 and CTSB (Figure 5C). Surprisingly, we observed decrease of PLD3 exonuclease activity in both sgSELENOO and sgFICD cells, when compared to parent Hap-1 cells, which suggests dysregulation of cellular response to CpG^O^-DNA stimulation (Figure S7A). In our previous work incubation of SH-SY5Y cells with monensin or bafilomycin A1 led to a strong increase in PLD3 AMPylation (Figure S9) (31). Surprisingly, monensin treatment did not result in an increase in PLD3 AMPylation in Hap1 cells, which further suggests a cell type specific regulatory mechanism of PLD3 AMPylation (Figure 5C). The complementary analysis of sgFICD cells without or with monensin treatment further showed the absence of AMPylation on PLD3 and HSPA5 (Figure 5E).

### FICD co-expression decreases PLD3 levels in HEK293T cells

FICD catalyzes both AMPylation and deAMPylation. Switching between the two functionalities is mediated by an autoinhibitory α-helix, which is contingent on the critical glutamate residue E234.^4^ To investigate whether the formation of different PLD3 forms is influenced by FICD activity, we co-expressed wt PLD3 and three variants of FICD – wt FICD, the highly active mutant FICD E234G, which lacks deAMPylation activity and the catalytically inactive mutant FICD H363A.^1^ First, the LC-MS/MS-based whole proteome analysis revealed that FICD E234G leads to significantly decreased levels of PLD3 compared to wt FICD and inactive FICD H363A (Figure 6A and Table S11). Complementary Western blot analysis confirmed the overall lower protein levels of PLD3 due to FICD E234G expression which rendered the soluble PLD3 levels difficult to detect (Figure 6B). This suggests that FICD might regulate PLD3 levels directly through AMPylation and indirectly by switching a yet unknown regulatory loop. Furthermore, expression of FICD E234G led to significant changes on whole proteome level (Figure 6C and Table S11). Interestingly, FICD E234G induced decrease in PLD3 levels was countered by an increase of HSPA5 and other UPR-associated proteins including HSPA6, HSB1, DNAJB1 DNAJB4, while no change was observed for the expression of housekeeping genes such as H3-2 (Figure 6D and Table S11). There is an overlap of five proteins which are upregulated when FICD E234G is overexpressed in comparison to all other conditions. This set of upregulated proteins include BAG3, HSPB1, HSPA6, DNAJB1 and DNAJB4 suggesting increasing cellular stress and induction of heat-shock proteins. Only coagulation factor V (F5) was significantly downregulated in FICD E234G cells comparing to other FICD mutants. The most dysregulated proteins were observed when comparing only AMPylating FICD E234G with catalytically inactive FICD H363A variant. From total of 181 dysregulated proteins, 152 proteins are upregulated including TPM3, CHP1, GLA, COPRS, NRAS, HRAS, B4GALT7 and COPRS (Figure 6C).

**Figure 6.**
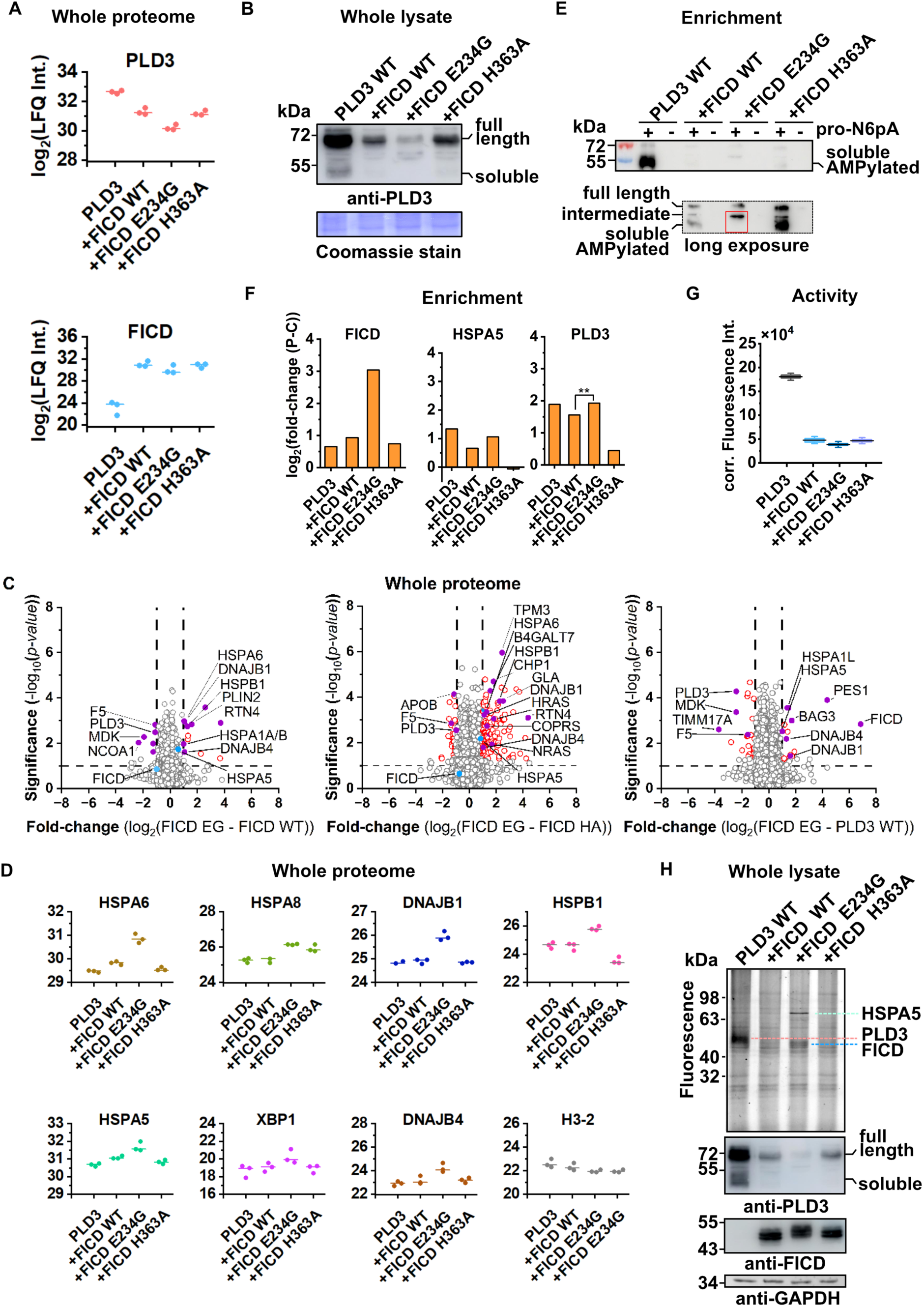
FICD-mediated AMPylation accelerates PLD3 degradation in HEK293T cells. **A**) Proteomics analysis of PLD3 levels changes in co-expression experiment of PLD3 with FICD variant. Of note, FICD was found and quantified in all replicates of FICD overexpressing cells, while only in one replicate from untreated cells. Representative result from two independent runs. Samples were prepared in triplicates (*n = 3)* from one protein lysate. **B**) Western blot using anti-PLD3 of whole cells lysate confirms decreasing amount of full-length and soluble PLD3 when co-expressed with FICD E234G. One representative blot of three independent experiments is shown. **C**) Volcano plots visualizing changes in protein expression on whole proteome level in HEK293T cells overexpressing FICD variants. Representative result from two independent runs. Samples were prepared in triplicates (*n = 3)* from one protein lysate. **D**) Profile plots of selected up- and down-regulated proteins upon FICD variants overexpression. **E**) Pro-N6pA-based enrichment of AMPylated proteins followed by Western blot using anti-PLD3. One representative blot of two independent experiments is shown. **F**) Pro-N6pA-based enrichment of AMPylated proteins followed LC-MS/MS analysis. The bar plots show the fold-enrichment of AMPylated proteins. Representative result from two independent runs. Samples were prepared in triplicates (*n = 3)* from one protein lysate. **G**) Box plot visualizing the drop of PLD3 exonuclease catalytic activity when co-expressed with FICD variants, *n* = 3 **H**) SDS-PAGE-based analysis of pro-N6pA reported AMPylation upon co-expression of PLD3 wt with FICD variants. One representative blot of two independent experiments is shown.

In opposite, the downregulated proteins include apolipoprotein B (APOB). Second, the analysis of PLD3 AMPylation status using the pro-N6pA probe upon co-expressions of FICD variants and subsequent Western blot with anti-PLD3 antibody showed a strong overall decrease in amount of AMPylated soluble PLD3 (Figure 6E). Surprisingly, in case of wt PLD3 together with FICD E234G co-expression, we observed strictly the intermediate form of PLD3 pointing towards malfunctioning downstream processing of PLD3 caused by missing deAMPylation activity (Figure 6E). Furthermore, the MS-based analysis of enriched AMPylated proteins showed significant FICD self-AMPylation and thus confirming the previously described *in vitro* observations (Figure 6F and Table S12) (53). Analysis of PLD3 AMPylation shows a significant increase in fold-enrichment between FICD wt and FICD E234G (Figure 6F). The normalization to the total and soluble form of PLD3 suggests that FICD might indeed catalyze AMP-transfer to PLD3 (Figure 6B and 6F). Of note, the MS-based analysis determines only the relative AMPylation ratio as it compares control to pro-N6pA enriched samples. Third, the PLD3 exonuclease activity has dropped accordingly correlating with the levels of soluble PLD3 (Figure 6G). The activity drops to the similar levels as for PLD3 point mutants lacking the AMPylation sites (Figure 2G). Finally, the in-gel analysis of pro-N6pA labelling confirmed the low amounts of soluble PLD3 when co-expressed with FICD variants and corresponding disappearance of fluorescent signal of soluble PLD3, which was visible only for wt PLD3 (Figure 6H). Together, the FICD AMPylation activity leads to decrease of both full-length and soluble PLD3 forms. The FICD H363A lacking deAMPylation activity obstruct the formation of soluble PLD3 and yielded only the intermediate PLD3 form.

### Full-length PLD3 is the major constituent of lysosomes

FICD is localized to the ER, while the PLD3 soluble catalytically active domain was previously described to be present in lysosomes, where it is transported through the endosomal pathway (39). However, the above-described interaction between PLD3 and FICD suggest that PLD3 mainly interacts with proteins localized to the ER and Golgi (Figure 4F). This raised the question, in which subcellular compartment AMPylation and deAMPylation are taking place and what the major form of PLD3 in lysosomes is. To this end, we used LysoIP with the transient overexpression of lysosomal TRPML1 Ca^2+^ channel bearing a triple human influenza hemagglutinin (HA) tag (Figure 7A) (54). The efficient enrichment of the lysosomal fraction was determined by immunoblotting, using LAMP1 as a lysosomal marker, and LC-MS/MS proteomics, both confirming the successful isolation of lysosomal fraction (Figure 7B, 7C and Figure S12). Surprisingly, we observed solely the full-length PLD3 form present in the enriched lysosomal fraction, while both the soluble and full-length PLD3 forms were clearly observed in original lysates before the LysoIP (Figure 7C). Together, LysoIP suggests that PLD3 in lysosomes is present in its full-length and not its soluble form as thought previously.

**Figure 7.**
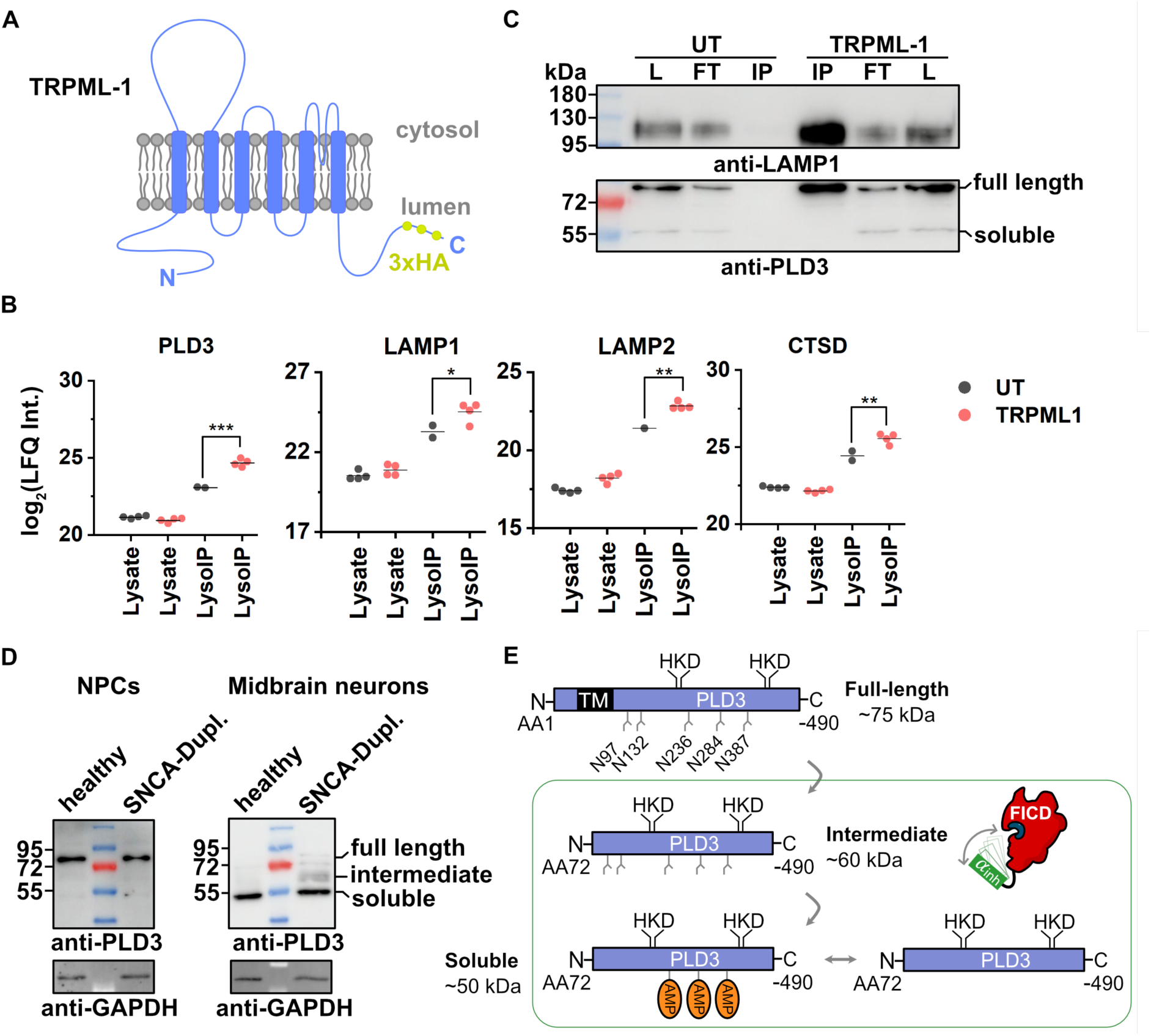
LysoIP renders full-length PLD3 as the main constituent of lysosomes. **A**) Scheme of TRPML1-3xHA construct. TRPML1 consists of six transmembrane helices and an extracytosolic/luminal domain, the hallmark of the TRPML family (65). The 3xHA tag is fused to the disordered C-terminus. **B**) Profile plots of PLD3 and lysosomal markers obtained from LC-MS/MS analysis of lysate and after lysosomal IP. UT-untreated or TRPML1-3xHA tag containing HEK293T cells. LysoIP is an enrichment experiment comparing protein background binding from untreated cells with lysates from TRPML1-3xHA containing cells. Of note, the protein of interest, such as PLD3, was found and quantified in all replicates from TRPML1-HA containing cells, while only in two or less replicates from untreated cells. One replicate represents an individual LysoIP sample. **C**) Western blot with anti-PLD3 of whole lysate (L), flow through (FT) and lysosomal IP (IP) fraction for TRPML1 overexpressing or untreated (UT) HEK293T cells. One representative blot from two independent LysoIP experiments is shown. **D**) Western blotting of PLD3 in NPCs from a healthy donor compared to midbrain neurons from a PD patient with SNCA duplication. In PD-derived neurons higher levels of PLD3 intermediate forms were observed. **E**) Proposed post-translational processing of PLD3 resulting in three different forms. In green rectangle are PLD3 forms regulated by FICD AMPylation and deAMPylation activity, which is governed by its inhibition α-helix.

### PLD3 forms are altered in PD patient-derived neurons

The function of PLD3 has been previously linked to neurodegenerative conditions, such as AD (55). Western blot analysis of PLD3 revealed association of distinct PLD3 forms with differentiated neurons (see Figure 1F). In human iPSCs (hiPSCs), the predominant form of PLD3 is the full-length version. However, in neurons, the soluble form of PLD3 predominates, with more than 100-fold higher levels compared to HEK293T cells and over 17-fold higher levels than in SH-SY5Y cells (Figures 1F and S2).

To investigate the relationship between different forms of PLD3 and neurodegenerating neurons, we characterized PLD3 in hiPSC-derived NPCs and midbrain neurons from a PD patient carrying a heterozygous duplication of the SNCA gene, which codes for α-synuclein, and compared them to neurons from a healthy donor. Our previous study demonstrated that midbrain neurons carrying SNCA duplication exhibited neurite deficits (45), a pathological event that precedes neuronal loss (56).

Confirming our observation in hiPSC and neurons (Figure 1F), the main forms in proliferating NPCs was the full-length form. The predominant form in differentiated neurons is soluble PLD3 (Figure 7D). Interestingly, in PD patient-derived midbrain neurons, we observed the presence of intermediate and full length PLD3, when compared to control neurons, suggesting a delayed or impaired processing of PLD3.

## Discussion

Protein PTMs can swiftly regulate protein catalytic activity, protein-protein interactions, and protein subcellular localization. Therefore, PTMs provide the cell with an efficient and fast mechanism to adjust metabolic pathways by reacting to changes in the intracellular and extracellular environment. Indeed, protein PTMs have been shown to play an important role in many biological processes, while their dysregulation can be often associated with a disease state (57, 58). The number and stoichiometry of many known protein PTMs differ dramatically in cell types, leading to the concept of proteoforms (58, 59). A proteoform is understood as an individual protein isoform and splice variant with defined PTM status. The different proteoforms resulting from expression of a gene may have different subcellular localization, catalytic activity, and interactions with other proteins. Thus far, multiple protein PTMs have been closely associated with neurodegenerative diseases for example by influencing aggregation properties of Tau and α-synuclein (60, 61). The first step to deconvolute such complex network and dynamic changes of proteoforms resulting in the onset of neurodegeneration or reversing the metabolism towards the homeostasis is understanding properties of the individual protein ‘players’ or proteoforms including plethora of ever emerging protein PTMs. Protein AMPylation is a reversible modification on the side-chain hydroxyl group of Tyr, Ser or Thr residues (62). In human cells, the most prominent AMPylated protein is HSPA5, the main activator of UPR. HSPA5 is AMPylated by FICD to inhibit its chaperone activity once the ER stress is released and elevated HSPA5 expression is decreasing to its basal level (11, 13, 20). In addition, the pool of AMPylated HSPA5 in the cells can provide quick access to active HSPA5 by FICD-catalyzed deAMPylation (21). The obstacle to address functionally FICD-related AMPylation/deAMPylation is its low abundancy in most cell lines, aforementioned dual catalytic activity and yet unclear mechanism(s) allowing FICD to switch between AMPylation and deAMPylation. Interestingly, a *Drosophila* FICD mutant lacking the deAMPylation activity was not viable and induction of excessive AMPylation by the a hyperactive FICD mutant led to scrambled eye development (10). Moreover, overexpression of FICD in developing cerebral cortex organoids revealed accelerated neuronal differentiation from NPCs, which resulted in the partial failure of newly produced neurons to migrate into the cortical layer (22). Another intriguing question concerns FICD co-substrate selectivity, as it seems highly likely that different nucleotide analogues serve as a substrate in living cells (29, 63). Furthermore, several dozen additional proteins were identified to be AMPylated, many of them lysosomal proteins known to be associated with neurodegeneration (18, 64).

The function of PLD3 is extensively studied in immune cells where it regulates the activation of TLR7 and TLR9 by cleaving ssRNA and ssDNA substrates (34, 40, 42, 66). Based on a GWAS study, PLD3 was linked to AD, but the putative mechanism by which PLD3 contributes to the pathology remains controversial. Studies with neuroblastoma cells identified mitochondrial DNA (mtDNA) as the major physiological source of PLD3 DNA substrate (35, 36, 67). Recently, an AD mouse model showed that lack of PLD3 activity improves axonal fitness and hence neuronal networking (33). While neurons are not immunologically active, the signaling pathways may be employed for other function as suggested for excitatory CA1 neurons to employ TLR9 inflammatory signaling to establish features characteristic for memory assemblies (68). As shown by us here and previously, there is a striking difference of PLD3 levels and proteoforms between immune and neuronal cells (31). The major PLD3 form in neurons is the proteolytically cleaved soluble PLD3 with highest AMPylation stoichiometry in mature neurons. Pull-down of pro-N6pA modified PLD3 shows that only the proteolytically processed soluble PLD3 form is AMPylated. There are several cognate questions remaining to answer about PLD3 function and processing in general and specifically in neurons, which might be important for understanding the mechanism relevant for the role in neurodegeneration.

In this work, we focused on the characterization of PLD3 AMPylation and its functional implications. We provide several lines of evidence showing that AMPylation is necessary for PLD3 proteolytic processing and 5’-3’ exonuclease activity. First, we established a mass spectrometry-based proteomics strategy for identification of PLD3 AMPylation sites, which led to identification of three AMPylation sites Y323, S365 and S380, all located in the soluble domain of PLD3. (Figure 2). The respective PLD3 point mutants resulted in complete loss or strong reduction of pro-N6pA labelling for S365A and S380A or Y323A and triple mutant, respectively. This led to overall lower levels of soluble PLD3 and markedly reduced exonuclease activity (Figure 2). Interestingly, mutation of Y323 to alanine not only led to a complete loss of AMPylation, but also restricted formation of soluble PLD3, and inhibited its catalytic activity. In contrast, the mutation of Y323 to phenylalanine partially restored AMPylation, presumably on the other sites, and resulted in increased levels of soluble PLD3, suggesting its critical role in processing (Figure 2). Therefore, simultaneous PLD3 AMPylation on multiple sites might possess a rheostatic function. Second, in immune pDC-like PLD3^-/-^ Cal-1 cells, TLR9 response was reversed by wt PLD3, but not by Y323A and the triple (S365A, S380A, Y323A) mutant, which suggests a rheostatic role of AMPylation and privileged function of AMPylation on Y323. PLD3 S365A and S380A mutants still led to partial processing of PLD3, which explains their catalytic activity in Cal-1 cells. Presence of both PLD4 and PLD3 in Cal-1 cells further underline cell type specific processing of ssDNAs and ssRNAs, and points toward distinct PLD3 function in immune and neuronal cells.

Third, immunoprecipitation of PLD3-FLAG significantly enriched FICD pointing towards their direct interaction. Interestingly, the reverse experiment in which FLAG-FICD was pulled-down corroborated the interaction and additionally revealed that FICD interacts specifically with an intermediate form of PLD3 (Figure 4). Since the intermediate form is the least abundant form in all tested cell types, this observation suggests that FICD-PLD3 interaction initiates fast PLD3 scission to release soluble PLD3. Moreover, the complementary proteomics analysis confirmed the Western blot results and identified a group of 70 proteins interacting with both, PLD3 and FICD. Interestingly, all common interacting proteins strictly localize to the ER and Golgi, while PLD3 immunoprecipitation together with lysosomal proteins was not observed, pointing towards PLD3 major localization and function in ER, endosomes and during the secretion process (Figure 4).

Fourth, we aimed to show whether co-expression of FICD E234G, which lacks the deAMPylation activity, together with PLD3 might lead to increased PLD3 AMPylation. Surprisingly, although there is a significant increase in PLD3 AMPylation, the comparison of FICD E234G to FICD wt and the catalytically inactive H363A variant shows a strong decrease in overall PLD3 levels (Figure 6), as observed by Western blot analysis and corroborated by LC-MS/MS proteomics. Therefore, in HEK293T cells FICD-mediated PLD3 AMPylation leads to its accelerated degradation. The whole proteome analysis suggests a strong upregulation of protein folding capacity, and UPR activation. Next, utilizing the pro-N6pA probe for enrichment of the AMPylated PLD3 and subsequent Western blot analysis shows that the co-expression of PLD3 with FICD wt and H363A variant leads to enrichment of mainly soluble PLD3 form, small amount of full-length and intermediate form. However, in case of FICD E234G no soluble PLD3 form could be detected. This further suggests that both AMPylation and deAMPylation activity of FICD are needed to properly process PLD3.

Fifth, lysosomes were previously identified as the major location of soluble and catalytically active PLD3 (39). However, we found only the full-length PLD3 form in lysosomes, while both the full-length and soluble form could be detected in whole cell lysates (Figure 7). Together with the PLD3 IP experiments, these results suggest that major subcellular localization and function of the PLD3 is in the ER and endosomes, where it might first regulate TLR9 signaling, before their relocalization to lysosomes.

Sixth, the possible dysregulation of PLD3 processing in neurodegeneration was explored in patient derived hiPSCs and midbrain dopaminergic neurons (Figure 7). The patient possessed known PD risk factor of α-synuclein gene duplication. The Western blot analysis showed hampered PLD3 processing with more abundant mixture of full-length and intermediate form in comparison to the healthy control, which may result in changes of downstream TLR9 signaling (Figure 7).

In summary, our study identifies three AMPylation sites in PLD3 and establishes AMPylation as a critical processing step required for production of catalytically active soluble PLD3 (Figure 7E). The direct interaction between PLD3 and FICD, together with co-expression experiments, suggests that FICD mediates AMPylation of PLD3 in living cells. In combination, accumulation of soluble PLD3 in neurons and observed functional role in neurodegenerative diseases renders AMPylation a plausible mechanism, which tightly regulates downstream TLR9 signaling. The identification if this regulatory cascade may offer desired pharmacological opportunities to improve neuronal fitness in neurodegeneration.

## Acknowledgement

This work was supported by the Deutsche Forschungsgemeinschaft (DFG, German Research Foundation) SFB1309 – 325871075 (PK), project 470553481 (LTJ), SFB1507 – 450648163 (UAH), SFB1278 – 316213987 (UAH) and EXC 2051 – project 390713860 (UAH), Boehringer Ingelheim Foundation – Plus 3 Program from (PK) and Exploration Grant (UAH), Liebig Fellowship from Verband der Chemischen Industrie (PK), European Research Council project StG 804182 (LTJ), Interdisciplinary Center for Clinical Research (IZKF, ELAN) P128 (WX), Johanna and Frieda Marohn-Stiftung, Pyroglutamic-acid-α synuclein (WX).

## Data availability

All MS data are available via ProteomeXchange with identifier PXD057707. Token: bKP4d9adfVvj

## Supplemental data

This article contains supplemental data.

