## Supplemental data for "AMPylation regulates PLD3 processing"

### Supplementary Figures

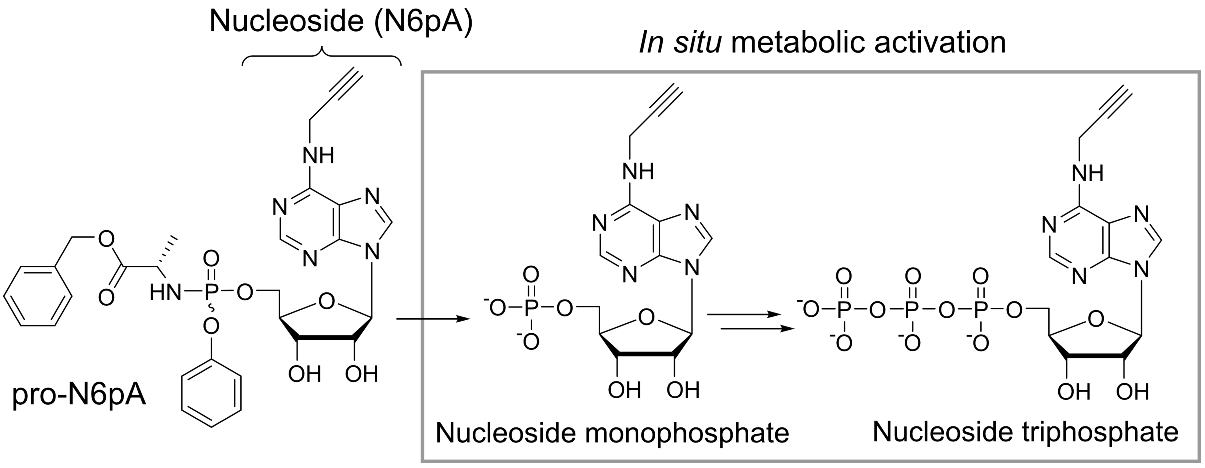

**Figure S1:** Scheme of the pronucleotide probe pro-N6pA and parent adenosine derivative (N6pA) and its in situ activation. The scheme was modified from that previously shown in ref.(1)

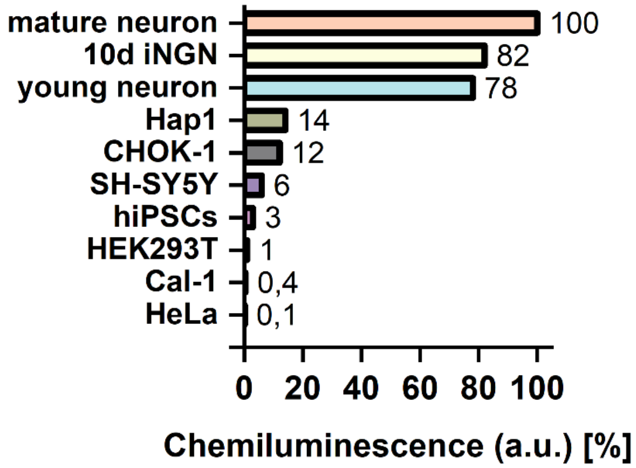

**Figure S2:** Quantification of PLD3 from western blot in Figure 1F. All PLD3 forms are accounted for. Protein amounts were normalized to anti-GAPDH loading control. Hap1 cells showing the highest levels of full-length PLD3 form. 17-fold more PLD3 is present in neurons in comparison to other cell types.

**S365 - unmodified peptide**

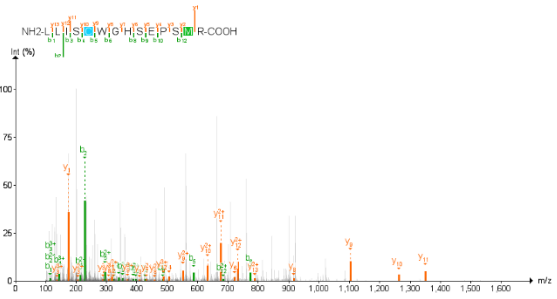

**S365 - modified peptide**

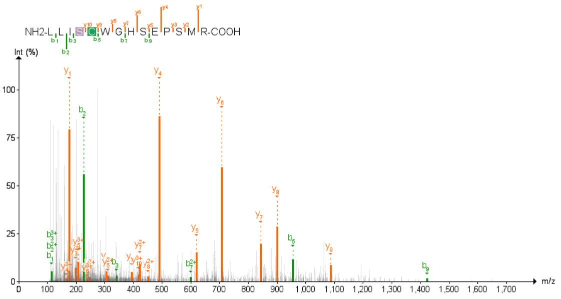

**Y323 - unmodified peptide**

**Y323 - modified peptide**

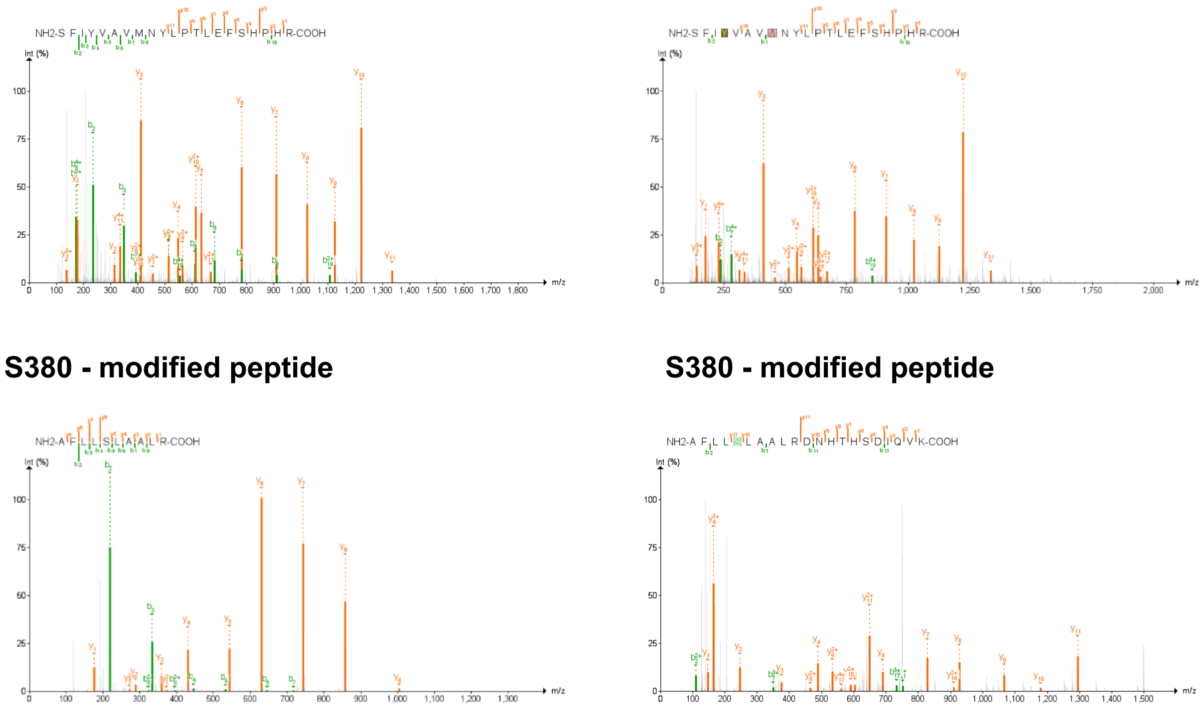

**Figure S3.** Representative assigned MS/MS spectra for identified peptides from PLD3 including those containing residues S365, Y323 and S380, which were found to be AMPylated.

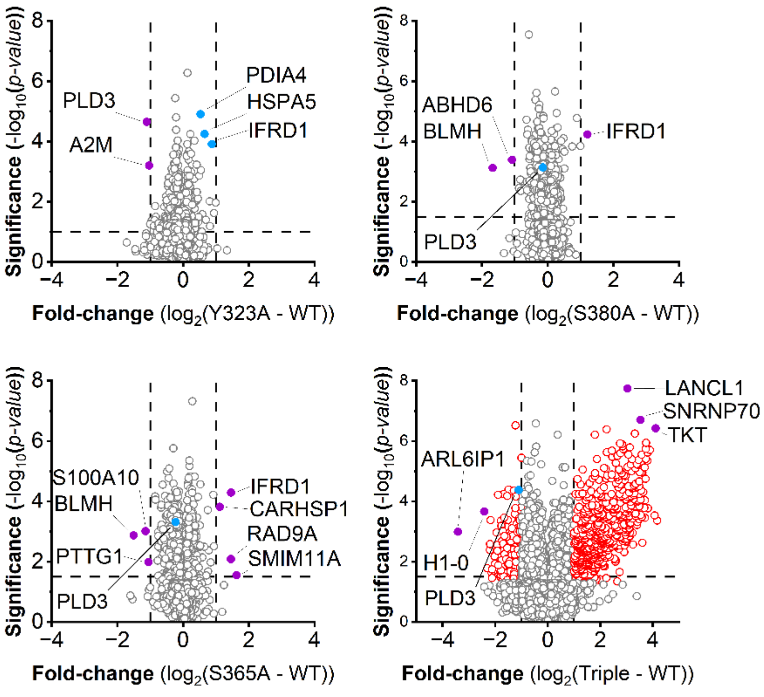

**Figure S4**. Whole proteome analysis of wt PLD3 and PLD3 Y323A, S365A, S380A and triple mutant in HEK293T cells.

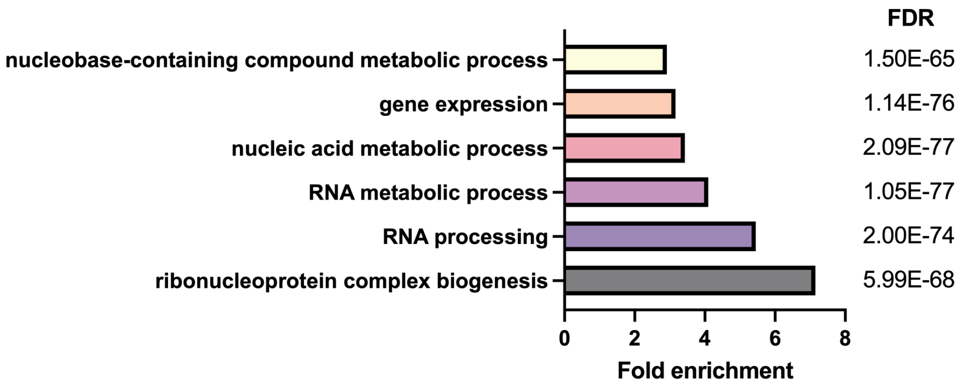

**Figure S5**. GO terms analysis of whole proteome changes (see Fig. S4) induced by the overexpression of triple (Y323A, S365A, S380A) PLD3 mutant in HEK293T cells. Both significantly up- and down-regulated proteins (see Figure S4) were used as the input for the analysis.(2)

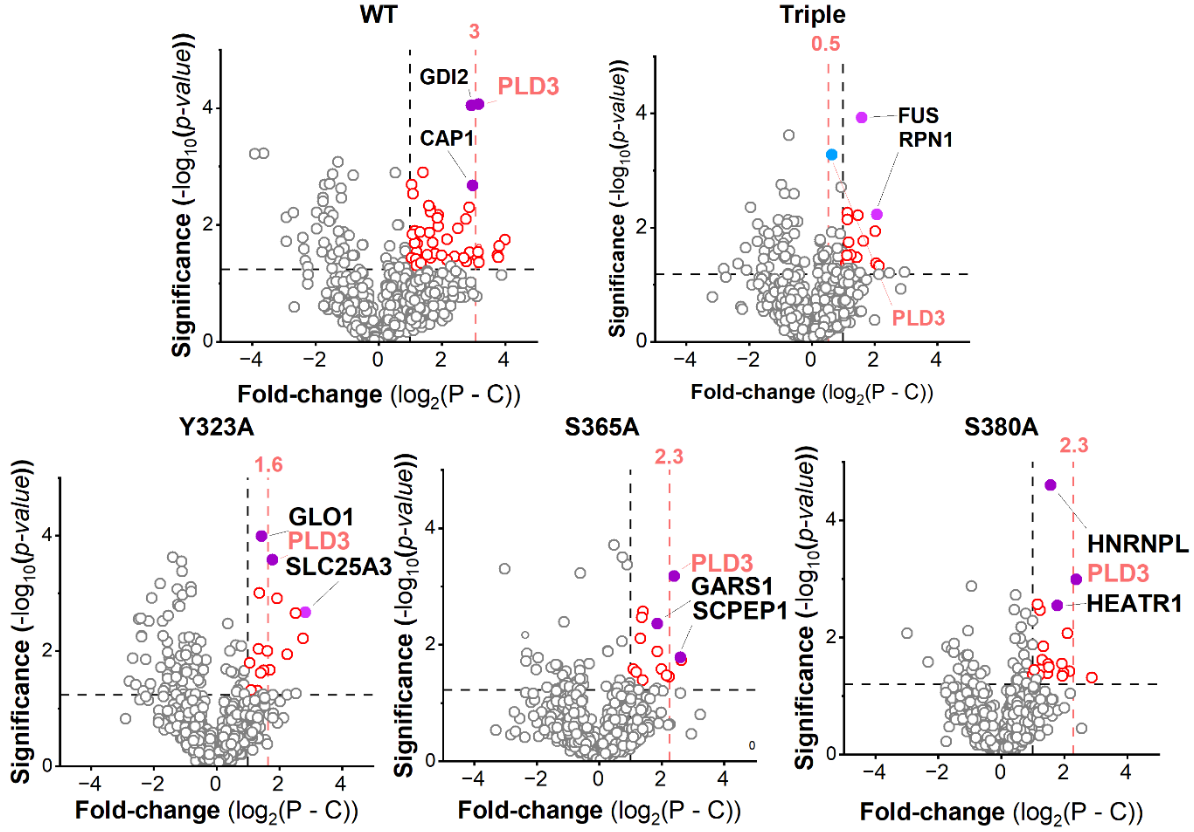

**Figure S6.** Enrichment analysis of wt PLD3, triple and single point mutants. P – pro-N6pA treated cells, C – control (DMSO treated), red circles – significantly enriched proteins, gray circles – not significantly enriched proteins.

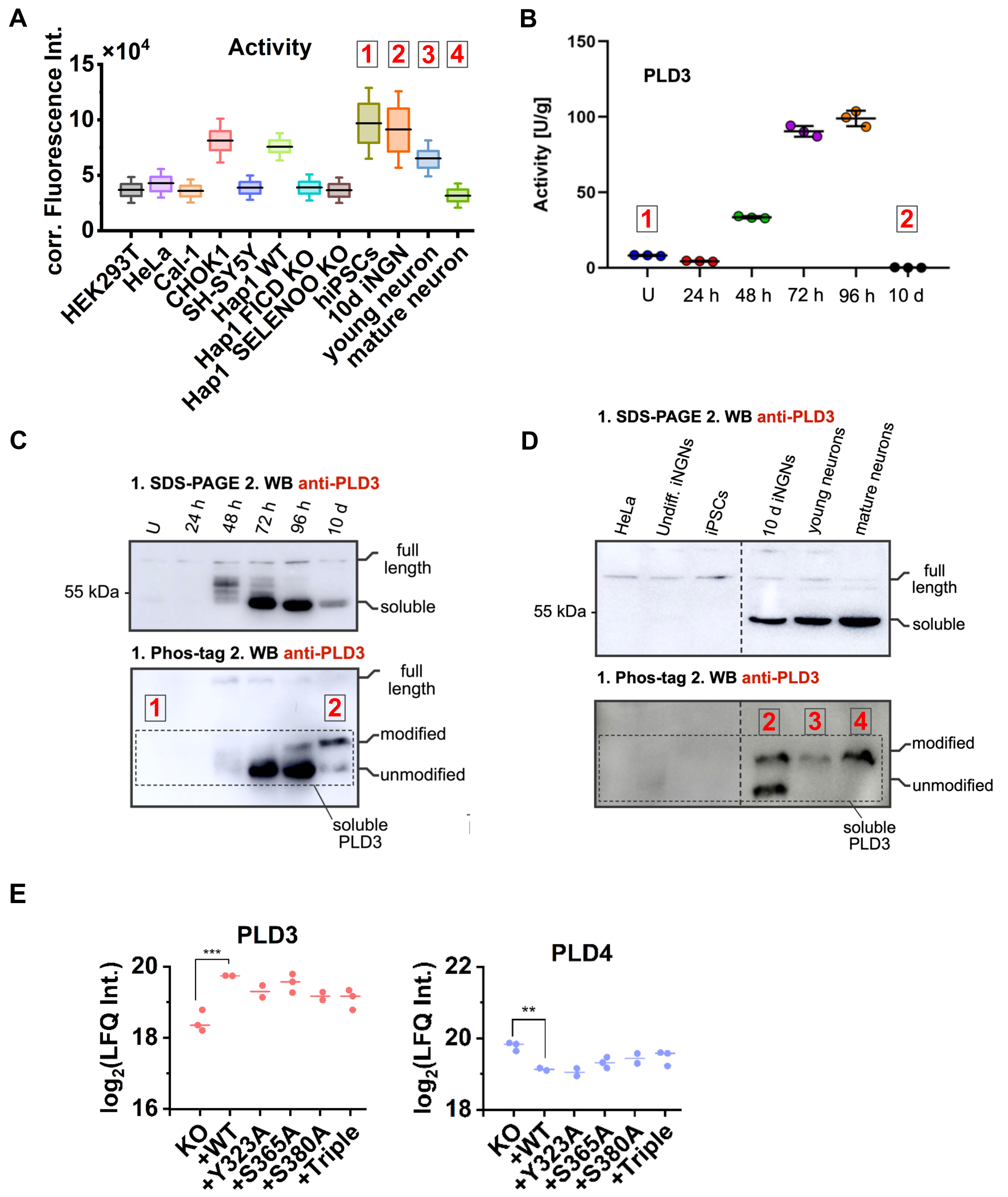

**Figure S7.** Analysis of PLD3 5' exonuclease activity and modification status.

(A) Box plots comparing PLD3 5'-3' exonuclease activity across different cell lines, *n = 3*.
(B) PLD3 5'-3' exonuclease activity during iNGNs differentiation to neurons (n = 3, line indicates the mean, error bars show the standard deviation). This figure has previously been published in ref. [9]
(C) Processing of PLD3 during iNGNs differentiation and maturation analysed by Phos-tag ligand SDS-PAGE separation and Western blotting using the anti-PLD3 antibody. Cropped region of a figure previously shown in ref. [9]
(D) SDS-PAGE and Phos-tag ligand SDS-PAGE separations followed by Western blot using the anti-PLD3 antibody show pronounced and nearly quantitative PLD3 modification. It shows comparison of

physiological dopaminergic neurons with iNGN forward reprogramming. Unmodified figure previously shown in ref. [9]

(E) Profile plots of PLD3 and PLD4 levels obtained from LC-MS/MS analysis of whole proteome analysis in PLD3-deficient Cal-1 cells overexpressing wt PLD3, its single mutants Y323A, S365A, S380A or the triple mutant.

iPSCs – induced pluripotent stem cells, U – undifferentiated iNGN stem cells. The numbers 1-4 in boxes link the exonuclease activity to PLD3 form visualized on Western blot in corresponding cell type.

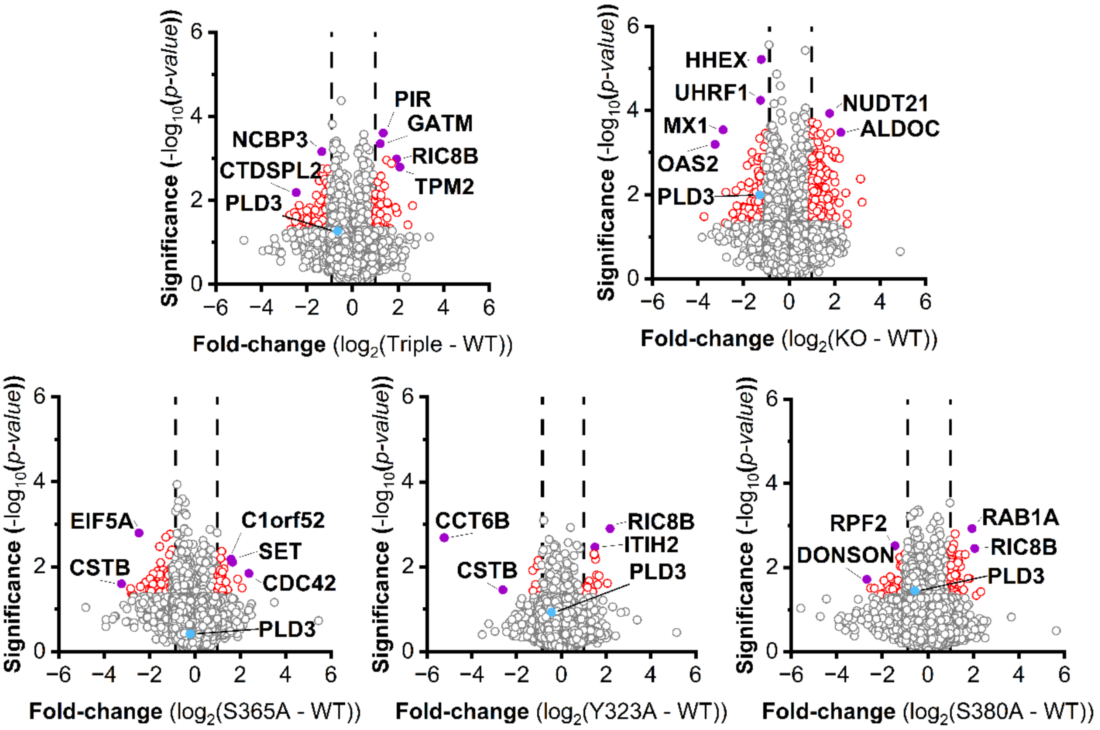

**Figure S8:** Whole proteome of PLD3-deficient Cal-1 cells stably expressing wt PLD3 and PLD3 single mutants Y323A, S365A, S380A or the corresponding triple mutant.

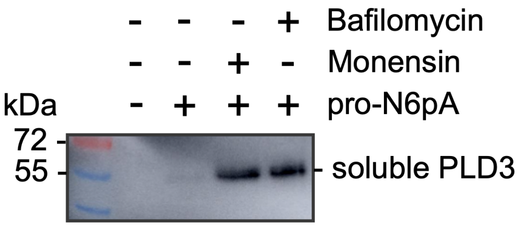

**Figure S9** Previously shown increase of PLD3 AMPylation on the soluble PLD3 in SH-SY5Y cells upon monensin or bafilomycin treatment. Western blot using anti-PLD3 antibody after enrichment of AMPylated proteins using pro-N6pA probe previously shown in ref.(3)

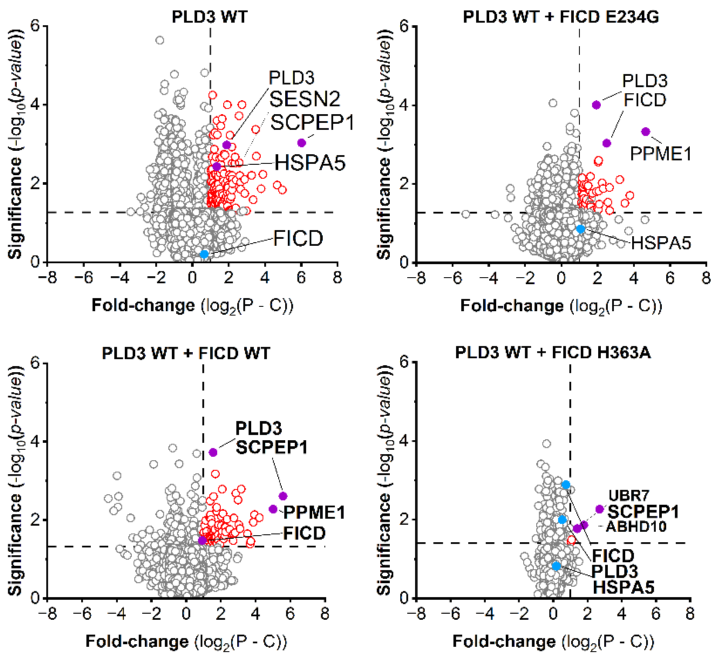

**Figure S10**. Enrichment analysis of wt PLD3 overexpression and co-expression of wt PLD3 together with wt FICD or E234G/H363A mutants in HEK293T cells. P – pro-N6pA treated cells, C – control (DMSO treated), red circles – significantly enriched proteins, gray circles – not significantly enriched proteins.

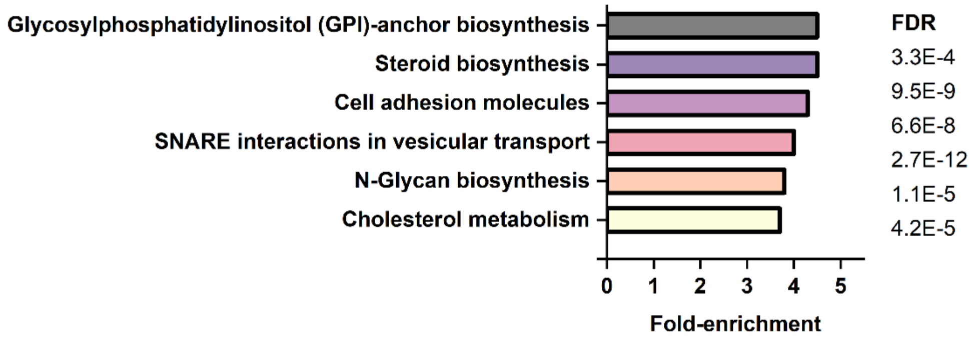

**Figure S11**. GO analysis of significantly enriched proteins after co-immunoprecipitation of PLD3-Flag from HEK293T cells.(2)

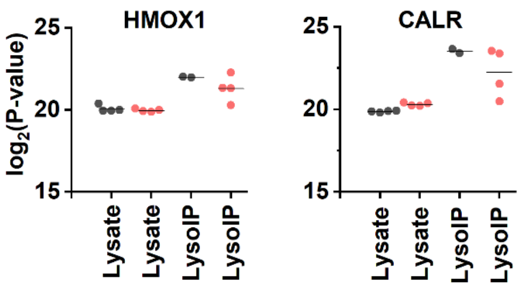

**Figure S12**. Profile plots of endoplasmic reticulum markers obtained from LC-MS/MS analysis of HEK293 cell lysate and after lysosomal IP.

**Table S1:** Sequences of oligonucleotides (5' to 3') used for cloning.

| Protein | Mutant | Sense | Sequence (5' to 3') | $T_a$ |
|---|---|---|---|---|
| PLD3 | Y323A | forward | GAGTTTCATCGCTGTCGCTGTCATGAAC TACCTGC | 66°C |
| | | reverse | CGGGCATTGTCCACCACG | |
| | S365A | forward | CCTGCTCATCGCTTGCTGGGGACACTC GGAG | 69°C |
| | | reverse | CGCACCTTGACGCCACGC | |
| | S380A | forward | CTTCCTGCTCGCTCTGGCTGCCC | 72°C |
| | | reverse | GCCCGCATGGATGGCTCC | |
| | *Δ*AA61 | forward | TGCTGCTCTGGGTTCCAGGTTCCACTGG TGACGAATACGGCGACTTG | 57°C |
| | | reverse | GTACCCATAGCAGCAGTGTGTCTGTTTC CATGCTAGCTGCTCGAAC | |
| FICD | H363A | forward | CGTTTACATCGCCCCTTTCATTGATG | 56°C |
| | | reverse | AGTTTATAATGGGCTAAGG | |
| | 3xFLAG | forward | CACGACATTGACTACAAGGATGATGACG ATAAGTAACCCTAGAAATCCTCAG | 58°C |
| | | reverse | GTCCTTATAGTCACCGTCATGATCTTTGT AATCGGGCTTCACAGGAAG | |

**Table S2:** Sequences of oligonucleotides (5' to 3') used for sequencing.

| Construct | Sense | Sequence (5' to 3') |
|---|---|---|
| **PLD3, full length** | forward | GTCCTGCATACCAAGTTCTG |
| **PLD3, *Δ*AA61** | forward | GTGTTAGACTAGTAAATTGTCCGC |
| **FICD** | forward | CATCATCCAGGCGGACTACT |

**Table S3:** Conditions for PCR cycle.

| Step | Temperature | Time | Cycle |
|---|---|---|---|
| **Denaturation** | 98°C | 30 sec | |
| **Annealing** | 98°C | 10 sec | 25 Cycles |
| | $T_a$ | 30 sec | |
| **Extension** | 72°C | 30sec/kb | |
| **final Extension** | 72°C | 2 min | |
| **Hold** | 4°C | | |

**Table S4:** Additional settings for AMPylation site-identification using DIA workflows offered by MSFragger.

| FragPipe settings | |
|---|---|
| **MS Fragger** | |
| **Mass offsets** | 0, 329.0525/367.0681 restricted to STY |
| **Peak matching** | Precursor Mass Tolerance ($MS^1$): 10 ppm<br>Fragment Mass Tolerance ($MS^2$): 10 ppm |
| **Variable Modifications** | 329.0525 on STY (natural AMP)<br>367.0681 on STY (pro-$N^6$pA probe)<br>15.995 on M (Oxidation) |

| | |
|---|---|
| **Fixed Modifications** | 57.02146 on C (Carbamidomethylation) |
| **Umpire** | |
| **mass defect filter** | disabled |
| **remove background** | disabled |

**Materials**

| Reagents/Resource | Reference/Source | Catalog Number |
|---|---|---|
| **Cell line** | | |
| Human: Hap1 (wt/FICD KO/SELENOO KO) | Prof. Dr. Lucas Jae | N/A |
| Human: HEK293T | Prof. Dr. Thomas Carell | N/A |
| **plasmid DNA** | | |
| pFUGW-hsPLD3_wt_3xFLAG | Prof. Dr. Veit Hornung | N/A |
| pcDNA3.1_hsPLD3_wt | Prof. Dr. Veit Hornung | N/A |
| pFUGW-hsPLD3_Y323A_3xFLAG | | N/A |
| pFUGW-hsPLD3_Y323F_3xFLAG | | N/A |
| pFUGW-hsPLD3_S365A_3xFLAG | | N/A |
| pFUGW-hsPLD3_S380A_3xFLAG | | N/A |
| pFUGW-hsPLD3_Triple_3xFLAG | | N/A |
| pFUGW-hsPLD3-soluble_IGK_3xFLAG | | N/A |
| pCMV-FICD-wt | Dr. Pavel Kielkowski | N/A |
| pCMV-FICD-WT_3xFLAG | | N/A |
| pCMV-FICD-E234G | Dr. Pavel Kielkowski | N/A |
| pCMV-FICD-E234G_3xFLAG | | N/A |
| pCMV-FICD-H363A | | N/A |
| pCMV-FICD-H363A_3xFLAG | | N/A |
| pcDNA3-hsTRPML1-3xHis | Prof. Dr. Ute Hellmich | N/A |
| **Proteins** | | |
| anti-FICD (rabbit) | | Cat# HPA021390 |
| anti-FLAG affinity gel | Sigma-Aldrich | Cat#A2220 |
| anti-GAPDH | Bio-Rad | Cat#12004168 |
| anti-PLD3 (rabbit) | Thermo Fisher | Cat# HPA012800 |
| EndoH | New England Biolabs | Cat# P0702S |
| goat-anti-rabbit HRP | Thermo Fisher | Cat# 31460 |
| **Oligonucleotides** | | |
| quenched FAM-ssDNA substrate | Biomers.net | N/A |
| **Chemicals and Enzymes** | | |
| Acetic acid (100%) | | |
| Acetone (HPLC grade) | VWR Chemicals | Cat# 20067.320 |
| Acetonitrile (LC-MS grade) | Thermo Fisher | Cat# A955-212 |

| | | |
|---|---|---|
| Agar-Agar | Carl Roth | Cat# 6494.3 |
| Alanyl-Glutamine | Sigma-Aldrich | Cat# G8541 |
| Ammonium Acetate | Sigma-Aldrich | Cat# 09689-1kg |
| Ammonium bicarbonate | Fluka | Cat# 09830-100g |
| Ammoniumperoxodisulfat | Sigma-Aldrich | Cat# 09913 |
| Ampicillin | Carl Roth | Cat# HP62.1 |
| Benchmark$^{TM}$ fluorescence Protein Standard | Invitrogen | Cat# LC5928 |
| Biotin-$PEG_3$-$N_3$ | Carbosynth | Cat# FA34890 |
| Bromphenol Blue | Fluka | Cat# 32768 |
| BSA | AppliChem | Cat# A6588 |
| Carboxylate-coated magnetic beads (hydrophobic) | Cytiva | Cat# 65152105050250 |
| Carboxylate-coated magnetic beads (hydrophilic) | Cytiva | Cat# 45152105050250 |
| Chloroacetamide | Sigma-Aldrich | Cat# C0267-100g |
| Color Prestained Protein Standard (10-250 kDa) | New England Biolabs | Cat# P7719S |
| cOmplete™ Protease Inhibitor Cocktail | Roche | Cat# 04693116001 |
| Coomassie Blue R-250 | Fluka | Cat# 27816 |
| $CuSO_4$ x 5 H2O | Acros | Cat# 10627162 |
| dd$H_2O$ (LC-MS grade) | Honeywell | Cat# 15665350 |
| DMEM (1x) | Sigma-Aldrich | Cat# D6546 |
| DMSO | Sigma-Aldrich | Cat# D4540 |
| DPBS (1x) | Sigma-Aldrich | Cat# D8357 |
| DTT | AppliChem | Cat# A2948,0025 |
| EDTA | BioChemica | Cat# A1103 |
| Ethanol (EtOH) | Merck | Cat# 34852 |
| Fetal bovine serum | Thermo Fisher | Cat# A3840001 |
| Fetal bovine serum (FBS) | Biochrom | Cat# S0115 |
| Fetal calf serum (FCS) | Thermo Fisher | Cat# 10270106 |
| Formic Acid (LC-MS grade) | Thermo Fisher | Cat# A117 |
| GelStain | Carl Roth | Cat# 3865.1 |
| GlutaMAX™ Supplement | Thermo Fisher | Cat# 35050038 |
| Glycerol | Sigma-Aldrich | Cat# G5516 |
| Hepes | Carl Roth | Cat# HN77.5 |
| HEPES solution | Sigma-Aldrich | Cat# H0887-100ML |
| IMDM (1x) | Sigma-Aldrich | Cat # I3390 |
| LB medium | Carl Roth | Cat# 66693 |
| MEM NEAA | Gibco$^{TM}$ | Cat# 11140035 |
| MES | Sigma-Aldrich | Cat# M3671 |
| Methanol (LC-MS grade) | Thermo Fisher | Cat# A456 |
| NP-40 | Sigma-Aldrich | Cat# 74385 |
| Opti-MEM$^{TM}$ I supplement | Thermo Fisher | Cat# 31985062 |
| Penicillin/Streptomycin | Gibco$^{TM}$ | Cat# 15-140-122 |
| Polyethylenimine (PEI) | Polyscience | Cat# 23966 |
| pro-N6pA | Ref. (1) | N/A |
| R848 (Resiquimod) | InvivoGen | Cat# tlrl-r848 |
| Recombinant Human IFN-γ | PeproTech | Cat#300-02 |
| Rotiphorese.Gel 30 (37,5:1) | Carl Roth | Cat# 3029.1 |

| | | |
|---|---|---|
| sodium dodecyl sulfate | AppliChem | Cat# A2572 |
| Sodium pyruvate | Thermo Fisher | Cat# 11360039 |
| Streptavidin magnetic beads | New England Biolabs | Cat# S1420S |
| TAMRA-$N_3$ | Baseclick | Cat# BCFA-008-1 |
| TBTA | TCI | Cat# T2993 |
| TCEP | Carbosynth | Cat# FT01756 |
| TEAB (1 M) | Sigma-Aldrich | Cat# T7408 |
| TEMED | Sigma-Aldrich | Cat# T9281 |
| Tris-base | Thermo Fisher | Cat# 10724344 |
| Triton X-100 | Sigma-Aldrich | Cat# X100 |
| Trypan Blue | Thermo Fisher | Cat# 11538886 |
| TrypLE Express | Thermo Fisher | Cat# 12604013 |
| Trypsin (sequencing-grade) | Promega | Cat# V5113 |
| Tween 20 | Sigma-Aldrich | Cat# P6585 |
| Urea | AppliChem | Cat# A1049 |
| **Software** | | |
| DIA-NN | Ref. (4) | github.com/vdemichev/DiaNN |
| FAIMS MzXML Generator | | github.com/coongroup/FAIMS-MzXML-Generator |
| FragPipe | Ref. (5) | fragpipe.nesvilab.org/ |
| FreeStyle | Thermo Fisher | N/A |
| MaxQuant | Ref. (6) | www.maxquant.org/download_ asset/ maxquant/latest |
| MSConvert | Ref. (7) | proteowizard.sourceforge.io/ download.html |
| Origin (9.9.5.171) | N/A | www.originlab.com/ |
| PDV viewer | Ref. (8) | github.com/wenbostar/PDV |
| Perseus (1.6.10.43) | Ref. (9) | maxquant.net/download_asset/ perseus/latest |
| **Kits and Other** | | |
| FAIMS Pro Duo Interface | Thermo Fisher | N/A |
| hIFN-beta DuoSet ELISA | Novus Biologicals | Cat# DY814-05 |
| Lumox mutliwell 96 well plate | Sarstedt | Cat# 94.6120.096 |
| Microlab Prep robot | Hamilton | N/A |
| Orbitrap Eclipse Tribrid Mass Spectrometer | Thermo Fisher | N/A |
| PEPMAP100 C18 5UM 0.3X5MM | Thermo Fisher Scientific | Cat# 160454 |
| PicoTip™ Emitter, Silica Tip™ | New Objectives | FS360-75-8-N-20-C15 |
| Pierce® BCA Protein Assay Kit | Thermo Fisher | Cat# 23225 |
| Plasmid Midiprep Kit | Zymo Research | Cat# D4200 |
| Plasmid Miniprep Kit | New England Biolabs | Cat# T1010S |
| Preparative HPLC | Agilent | N/A |
| ReproSil-Pur 120 C18-AQ, 1.9 µm | Dr. Maisch GmbH | Cat# r119.aq.0001 |
| ReproSil-Pur 120 NH2, 3 µm | Dr. Maisch GmbH | Cat# r13.a0.0001 |
| Sep-Pak C18 cartriges | Waters | Cat# 186000308 |
| Tissue Grinder | VWR | Cat# 432-0200 |

| | | |
|---|---|---|
| VP 250/10 Nucleodur 100-5 C18 | AppliChem | Cat# 762022.100RC |

### Uncropped SDS-gels and Western Blots

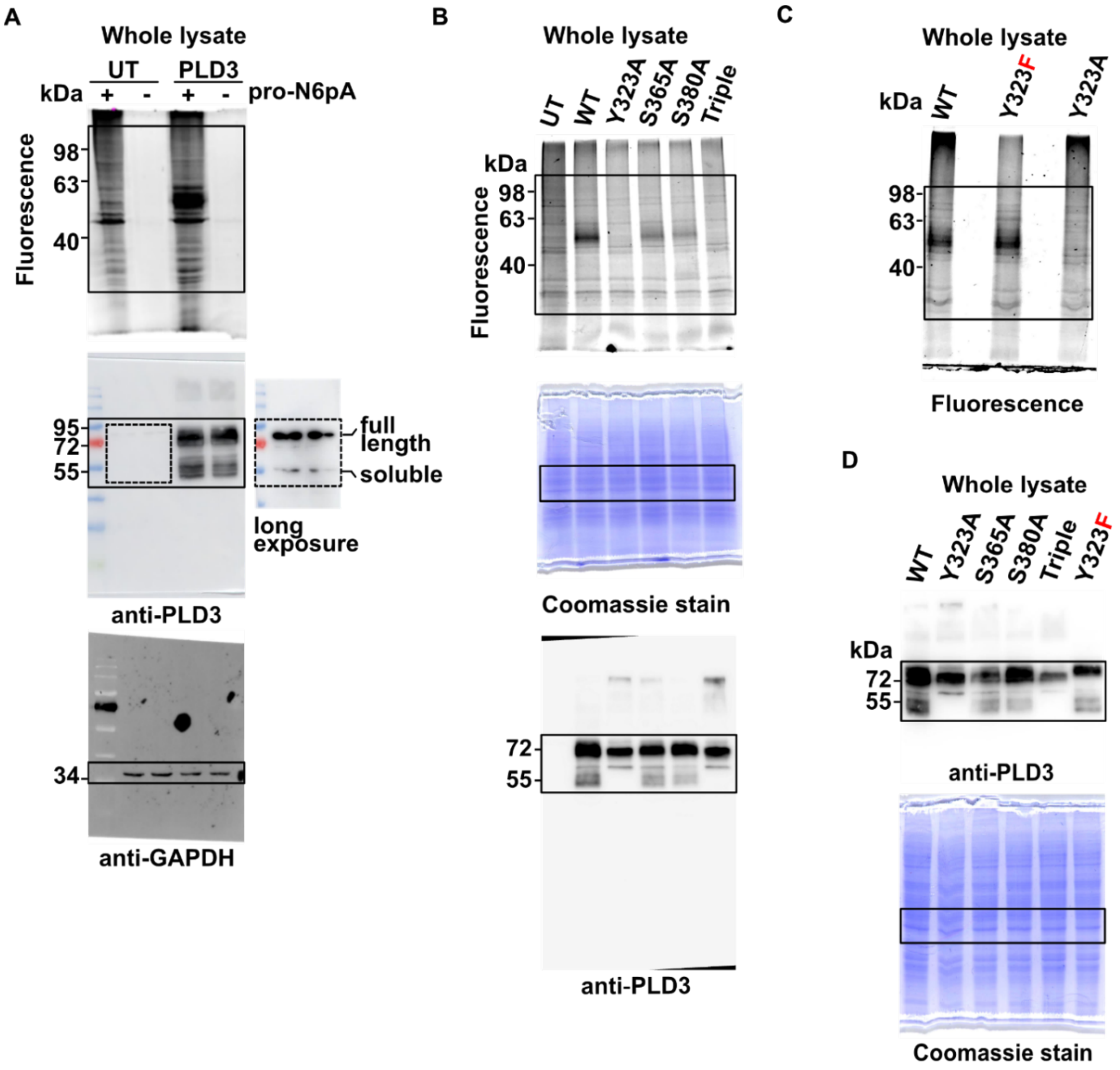

**Figure S13**: Uncropped gels shown in Figure 2.

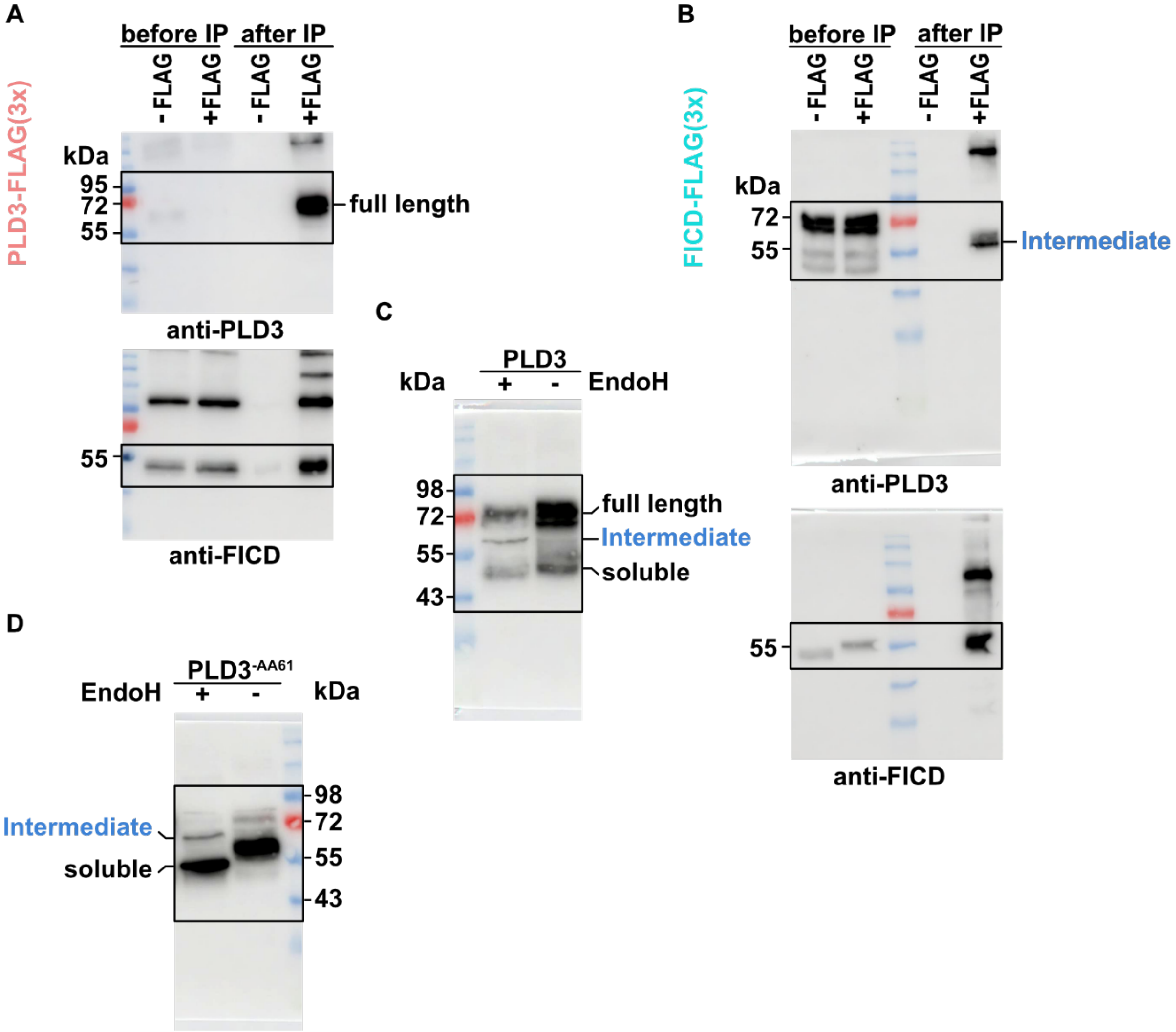

**Figure S14**: Uncropped gels shown in Figure 4.

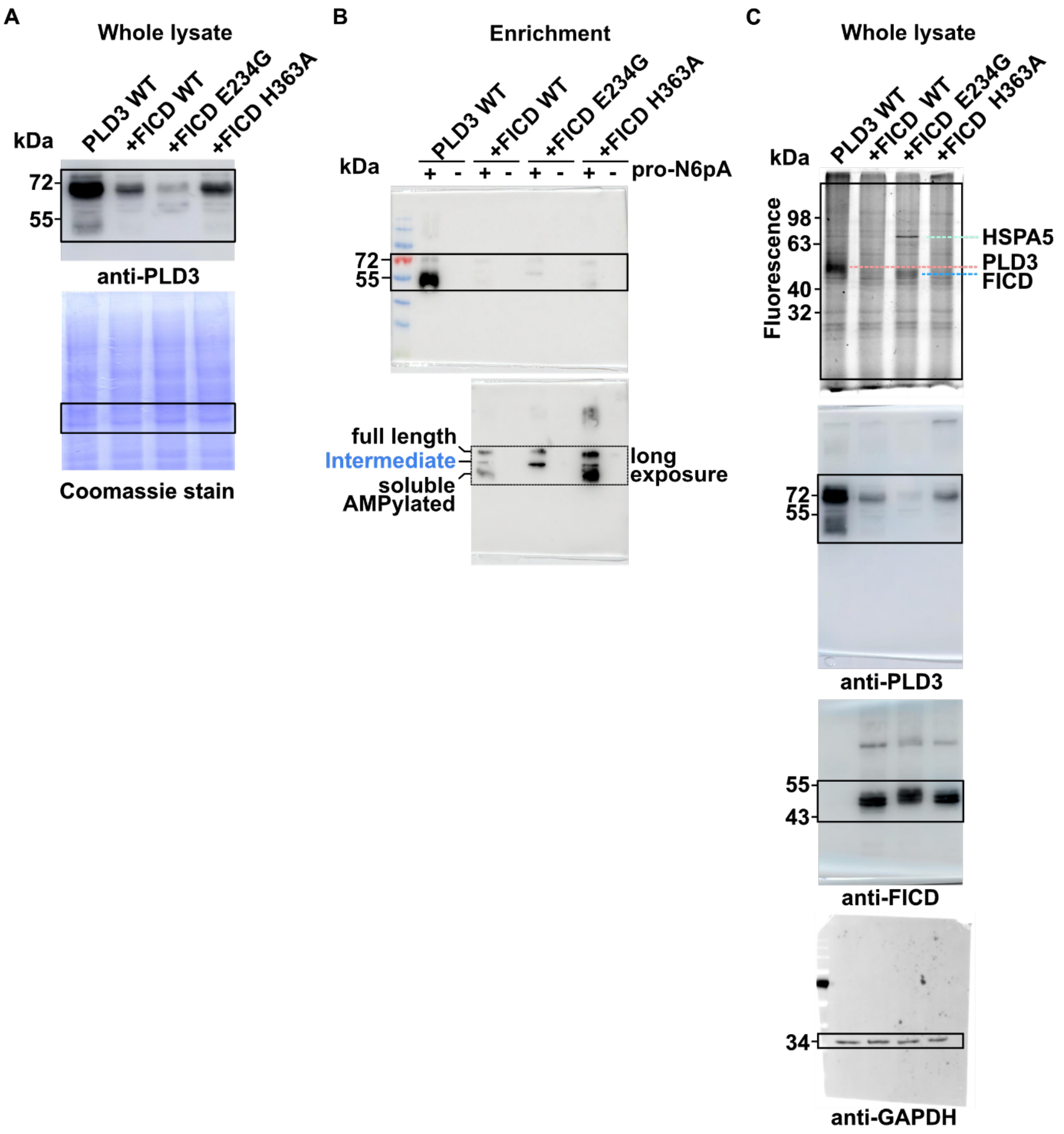

**Figure S15**: Uncropped gels shown in Figure 6.

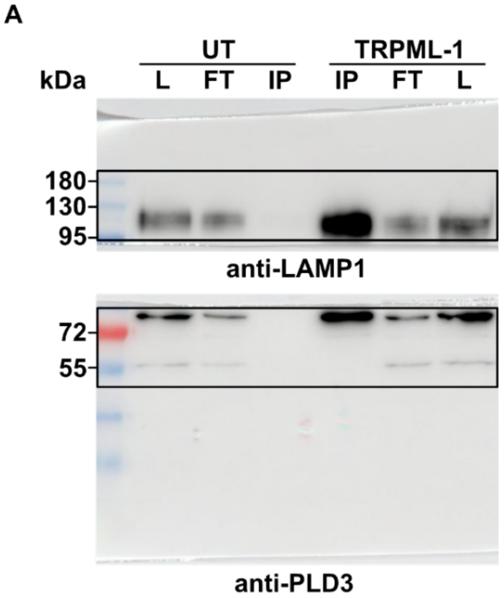

**Figure S16**: Uncropped gels shown in Figure 7.

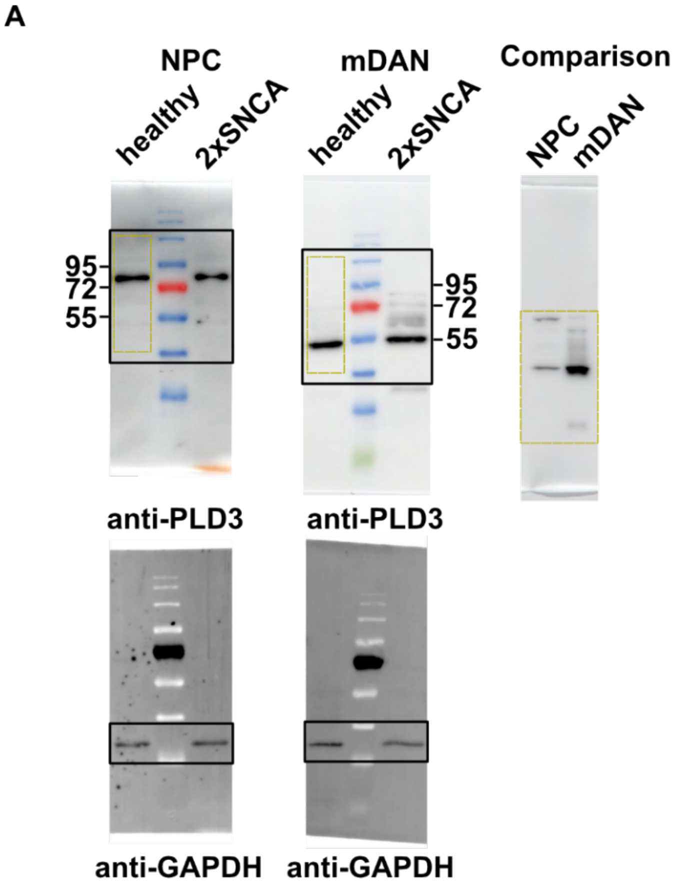

**Figure S17**: Uncropped gels shown in Figure 7.

1. Kielkowski, P., Buchsbaum, I. Y., Kirsch, V. C., Bach, N. C., Drukker, M., Cappello, S., and Sieber, S. A. (2020) FICD activity and AMPylation remodelling modulate human neurogenesis. *Nat Commun*. **11**, 517

2. Thomas, P. D., Ebert, D., Muruganujan, A., Mushayahama, T., Albou, L., and Mi, H. (2022) PANTHER: Making genome-scale phylogenetics accessible to all. *Protein Sci.* **31**, 8–22

3. Becker, T., Wiest, A., Telek, A., Bejko, D., Hoffmann-Röder, A., and Kielkowski, P. (2022) Transforming Chemical Proteomics Enrichment into a High-Throughput Method Using an SP2E Workflow. *Jacs Au*. 10.1021/jacsau.2c00284

4. Demichev, V., Messner, C. B., Vernardis, S. I., Lilley, K. S., and Ralser, M. (2020) DIA-NN: neural networks and interference correction enable deep proteome coverage in high throughput. *Nat. Methods*. **17**, 41–44

5. Kong, A. T., Leprevost, F. V., Avtonomov, D. M., Mellacheruvu, D., and Nesvizhskii, A. I. (2017) MSFragger: ultrafast and comprehensive peptide identification in mass spectrometry–based proteomics. *Nat. Methods*. **14**, 513–520

6. Cox, J., and Mann, M. (2008) MaxQuant enables high peptide identification rates, individualized p.p.b.-range mass accuracies and proteome-wide protein quantification. *Nat. Biotechnol.* **26**, 1367–1372

7. Chambers, M. C., Maclean, B., Burke, R., Amodei, D., Ruderman, D. L., Neumann, S., Gatto, L., Fischer, B., Pratt, B., Egertson, J., Hoff, K., Kessner, D., Tasman, N., Shulman, N., Frewen, B., Baker, T. A., Brusniak, M.-Y., Paulse, C., Creasy, D., Flashner, L., Kani, K., Moulding, C., Seymour, S. L., Nuwaysir, L. M., Lefebvre, B., Kuhlmann, F., Roark, J., Rainer, P., Detlev, S., Hemenway, T., Huhmer, A., Langridge, J., Connolly, B., Chadick, T., Holly, K., Eckels, J., Deutsch, E. W., Moritz, R. L., Katz, J. E., Agus, D. B., MacCoss, M., Tabb, D. L., and Mallick, P. (2012) A cross-platform toolkit for mass spectrometry and proteomics. *Nat. Biotechnol.* **30**, 918–920

8. Li, K., Vaudel, M., Zhang, B., Ren, Y., and Wen, B. (2018) PDV: an integrative proteomics data viewer. *Bioinformatics*. **35**, 1249–1251

9. Tyanova, S., Temu, T., Sinitcyn, P., Carlson, A., Hein, M. Y., Geiger, T., Mann, M., and Cox, J. (2016) The Perseus computational platform for comprehensive analysis of (prote)omics data. *Nat. Methods*. **13**, 731–740
